# Multiplexed Subspaces Route Neural Activity Across Brain-wide Networks

**DOI:** 10.1101/2023.02.08.527772

**Authors:** Camden J. MacDowell, Alexandra Libby, Caroline I. Jahn, Sina Tafazoli, Timothy J. Buschman

## Abstract

Cognition is flexible. Behaviors can change on a moment-by-moment basis. Such flexibility is thought to rely on the brain’s ability to route information through different networks of brain regions in order to support different cognitive computations. However, the mechanisms that determine which network of brain regions is engaged are unknown. To address this, we combined cortex-wide calcium imaging with high-density electrophysiological recordings in eight cortical and subcortical regions of mice. Different dimensions within the population activity of each brain region were functionally connected with different cortex-wide ‘subspace networks’ of regions. These subspace networks were multiplexed, allowing a brain region to simultaneously interact with multiple independent, yet overlapping, networks. Alignment of neural activity within a region to a specific subspace network dimension predicted how neural activity propagated between regions. Thus, changing the geometry of the neural representation within a brain region could be a mechanism to selectively engage different brain-wide networks to support cognitive flexibility.

## Main

Cognition arises from the dynamic interaction of brain regions^1,2^. Broad networks of regions are involved in sensory processing^9^, decision making^8^, and motor actions^10,11^. Which network is engaged changes over time, both during spontaneous behavior^3–6^ and in a task-dependent manner^2,8,12–14^. Routing neural representations through different networks is thought to support cognitive flexibility by changing how information is processed by the brain^15^. Several mechanisms have been proposed for determining how neural activity flows between regions^16^, including changes in the gain of neural response^15^ or synchrony of brain regions^17,18^. Yet, the mechanisms for engaging broader networks of regions remain unclear. Here we show interactions between regions are multiplexed, with different dimensions of neural activity within a region engaging different subspace networks. Aligning neural representations to one of these dimensions propagated the representation to the associated subspace network.

Using four Neuropixels probes^19^, we recorded neural activity in eight cortical and subcortical brain regions simultaneously (Fig. 1a-d; see methods). Overall, >6,500 neurons were recorded in hippocampus (HPC), thalamus (TH), prelimbic cortex (PL), frontal motor regions (FMR), retrosplenial cortex (RSP), visual-associated cortex (VIS), primary somatosensory cortex (SS), and whisker somatosensory cortex (WHS; see methods for detailed parcellation). All recordings were done while mice spontaneously engaged in different behaviors, ensuring a wide variety of cortex-wide neural dynamics^4,6^ (Fig. S1).

**Figure 1.**
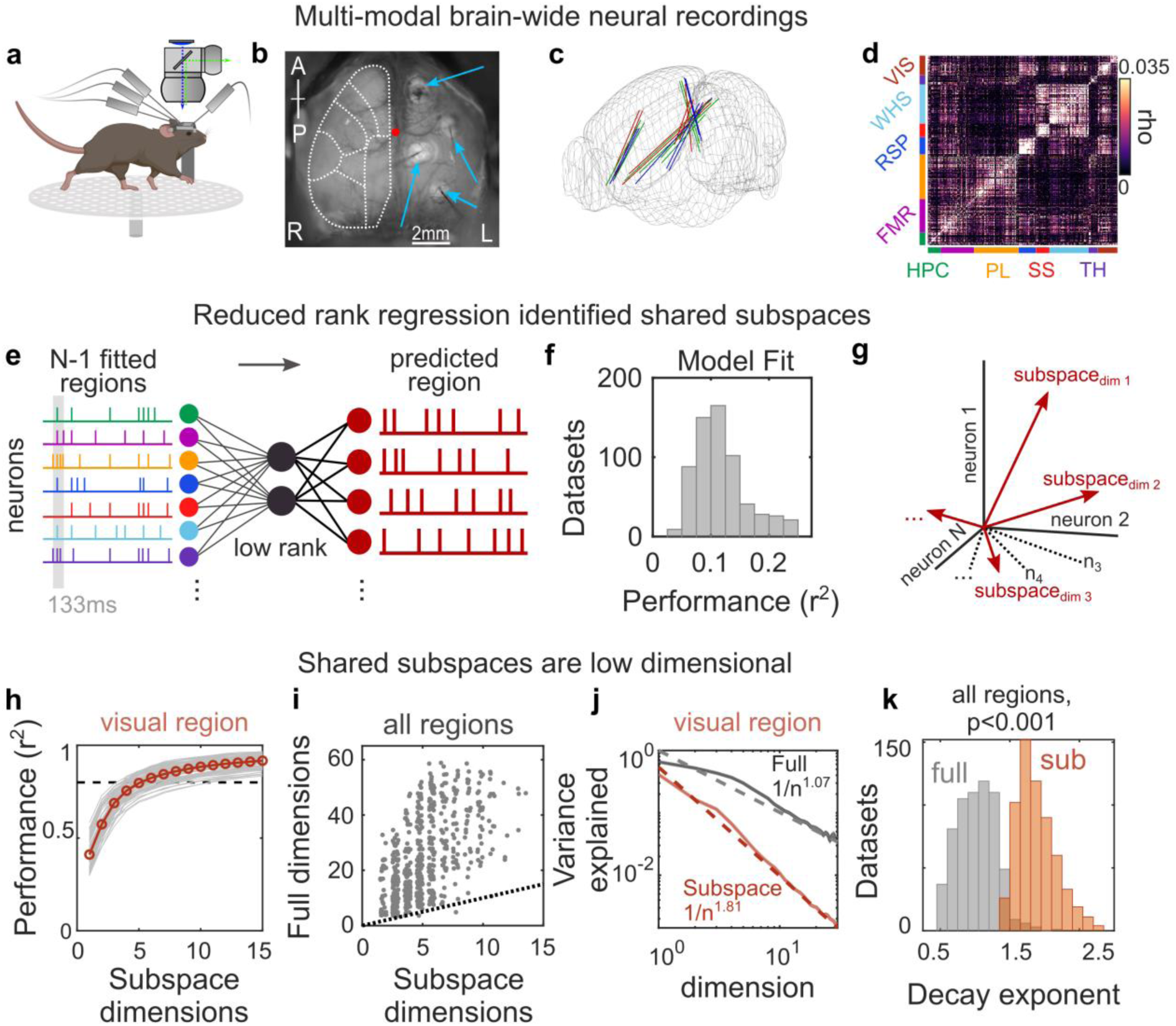
Interactions between brain regions are organized into subspaces. **(a)** Schematic of experiments combining neural recordings from four Neuropixels probes across cortical and subcortical regions and simultaneous widefield calcium imaging of neural activity across dorsal cortex. **(b)** Field of view of widefield imaging showing the dorsal cortex and four craniotomies with implanted electrodes (thin gray shadows; indicated by blue arrows). Dotted white lines outline general cortical regions (see Fig. S1 for parcellation). R and L denote animal’s right and left. **(c)** Reconstructed probe locations from six recordings in three mice; colors indicate mice. **(d)** Correlogram showing correlation between pairs of cells within a brain region compared to pairs across regions for an example recording. Color bars along axes correspond to brain regions, held constant throughout figures. **(e)** Schematic of reduced rank regression to predict activity of each neural population. Spontaneous spiking activity was binned to 133ms windows, and the evoked response was removed (see methods). The moment-to-moment spiking variability of one region was predicted using spiking variability of all other regions. **(f)** Cross-validated predictive performance of all regression models (models fit to n=630 datasets, see methods). **(g)** Schematic of the shared subspace, showing that the subspace is spanned by a set of dimensions within the neural activity of the predicted region that are correlated with neural activity in other regions. **(h-i)** A small subspace of neural activity within a region was shared with other regions. **(h)** Gray lines show cumulative percent of the explainable variance captured by each dimension of the reduced rank regression model (see methods). Red line shows mean performance. Gray lines show cross-validated performance from n=84 datasets. Black dotted line shows 80% of explainable variance. **(i)** Comparison between the number of subspace dimensions and the number of ‘local’ dimensions needed to explain 80% of the explainable variance within a region. Local dimensionality was estimated with a cross-validated measure^22^, which revealed that 24.1 dimensions (CI: 23.0-25.4) were needed to capture 80% of the explainable variance within each region. Dots show individual datasets (n=630). Dotted line shows unity. **(j)** Example results from visual cortex comparing the decay in the variance explained by each additional subspace or local dimension. Solid line and shaded regions show mean and SE, respectively (n=84 model fits). Dotted lines show fit of power law. **(k)** Histogram of power law exponents from fits to full (grey) and subspace (red) dimensionality across all regions (n=630 datasets).

To quantify how these eight brain regions were functionally connected, we used reduced rank regression^20^ to predict the pattern of neural activity in one brain region as a linear function of the activity in all other regions (Fig. 1e; note: models were fit to variance in neural activity, after removing evoked responses, see methods for details). All eight brain regions were interconnected. The regression models explained a significant amount of the variance in neural activity in all eight regions (Fig. 1f cross-validated r^2^=0.12, CI: 0.11-0.12, p<0.001 versus shuffled controls, see methods). Furthermore, each region contributed to the prediction of other regions; excluding any region decreased the cross-validated performance of >99% of the models (mean decrease of 7.6% explained variance, CI: 7.3-7.9%; this was robust to changes in model parameters, Fig. S2).

Brain regions were connected through a ‘shared subspace’. Reduced rank regression identified the set of orthogonal dimensions of neural activity within each brain region that were influenced by other regions (Fig. 1g; 𝑌_𝑑𝑖𝑚=𝑖_ ∼ β_𝑖_𝑋, where 𝑋 and 𝑌 are the neural activity in the fitted and predicted regions, respectively, β are linear weights, and 𝑖 ∈ [1. . 𝑁] is the dimension, sorted by variance explained; see methods for details). Consistent with previous work^21,22^, only a few dimensions of the neural population within a region were influenced by other regions. For example, within visual cortex, 80.0% of the variance in neural activity that could be explained by other regions was captured in the first 5 dimensions and 88% was captured by 10 dimensions (Fig. 1h). A similar pattern was seen across all brain regions; ≥80% of the explainable variance was captured by an average of 5.5 dimensions (CI: 5.3-5.7) and 10 dimensions captured an average of 90.0% of the variance (CI: 89.3-90.1%). Together, these dimensions encapsulated the subspace of population activity within each region that was shared with other regions^21,23,24^. Note that, although different dimensions of the shared subspace were orthogonal to one another, they engaged the same population of neurons (i.e., there were not independent populations of neurons supporting each dimension, Fig. S3).

The shared subspace within each region was significantly smaller than the ‘local’ space of that region’s population activity. On average, 4.6 times more dimensions were needed to explain 80% of the total variance in neural activity within a region (Fig. 1i; i.e., the local dimensionality^6^, see methods for details). The variance explained by adding a dimension to the local space ^22^ and shared subspace decreased as a power law (Fig. 1j shows decay for visual cortex). However, the contribution of additional dimensions decayed more rapidly in the shared subspace than the local space (shared decay exponent was -1.81, CI: -1.77--1.86, local: -1.07, CI: -1.04--1.11). This difference was seen for all regions (Figs. 1k) and reflects the fact that only a portion of the space of neural activity within a region was shared with other regions. Interestingly, the more ‘integrative’ prelimbic, frontal motor, and retrosplenial regions had higher dimensionality, both locally and shared with other regions, while the more ‘sensory’ visual, somatosensory, and whisker regions had relatively lower dimensionality (Fig. S4).

## Subspace dimensions interact with a distributed network of brain regions

Next, we aimed to understand how shared subspaces integrated neural activity from other brain regions. One hypothesis is that each dimension of the shared subspace reflects the exclusive coupling between a pair of regions, essentially acting as a ‘channel’ between the two regions. Alternatively, each dimension of the subspace may be coupled to multiple brain regions, integrating their activity into a single representation. To discriminate these hypotheses, we examined the beta-weights for each subspace dimension (Fig. 2a). Consistent with subspace dimensions integrating activity from a distributed network, the neurons with the largest weights were distributed across brain areas. For example, the largest weights for the first subspace dimension of the visual region were from neurons in retrosplenial, hippocampal and somatosensory regions (Fig. 2b; neural activity was normalized to allow for direct comparison of weights, see methods). To quantify this, we measured the cumulative distribution of weights for each region and found that retrosplenial, hippocampal, and somatosensory regions contributed the largest weights for the first subspace dimension of visual cortex (Fig. 2c; area under the curve, AUC, was significantly greater than 0.5, all p<0.001, see methods). In the majority of our recorded regions, the first two subspace dimensions integrated neural activity from multiple other areas (57%, CI: 38%-69%, see methods).

**Figure 2.**
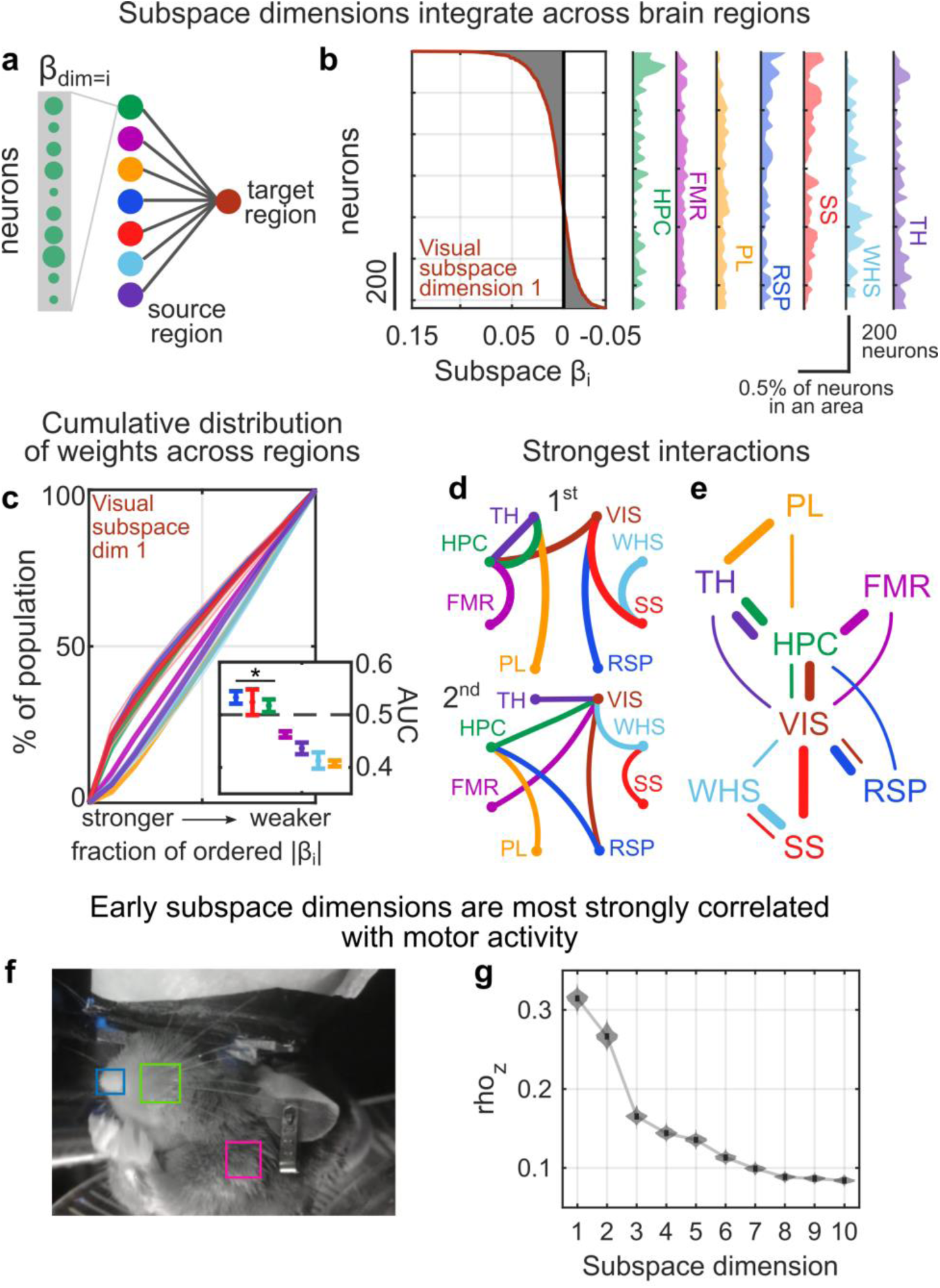
Subspaces between areas reflect brain-wide networks. **(a)** Contributions of neurons in a source region to predicting activity of a target region are given by their beta weights in the regression model. **(b)** Example distribution of beta weights (x-axis) across all neurons (y-axis) contributing to predicting activity along the first dimension of the shared subspace in visual cortex. Left plot shows distribution across all regions. Right plots show density histogram of weights within each region (normalized to account for different numbers of neurons). **(c)** Cumulative distribution of the percent of neurons within a source area contributing to the beta weights along the first dimension of the visual subspace (as in panel b). Weights are ordered from strongest to weakest. Lines show mean, shaded region shows bootstrapped 95% confidence interval from n=84 datasets (see methods). Inset shows the mean and confidence intervals of the area under the curve (AUC) for the contributions from each region. Asterix denotes significance at p<0.05 versus an AUC of 0.5, bootstrap test, n=1000 permutations. **(d)** Schematic showing the network of strongest (top) and second strongest (bottom) interactions between areas, across all datasets. Strength of contributions were taken as the AUC of beta weights (as in panel c, except averaged across all subspace dimensions). Colored lines indicate the subspace target region, i.e., the strongest contribution to FMR is from HPC (pink line). **(e)** Network graph showing the two strongest interactions between brain areas. Thicker/thinner lines show the first/second strongest interaction, respectively. **(f)** The relationship between subspace activity and motor activity was quantified by correlating the activity along each subspace dimension with the motion energy of animals’ nose (blue), whisker-pad (green), and shoulder (pink; see methods). **(g)** Bootstrapped distribution showing the average correlation (y-axis, z-transformed) between motor activity and activity along each subspace dimension (x-axis) across datasets. Subspace dimension 1 was significantly more correlated with motor activity than all other dimensions (p<0.001, n=1000 bootstraps from n=630 datasets).

In general, the hippocampus and visual cortex provided the strongest inputs to other regions. To quantify the strength of coupling between two regions, we measured the average AUC of the cumulative distribution of weights (as in Fig. 2c) across all subspace dimensions. Hippocampus and visual cortex were the first or second strongest influence on almost all of the other regions (Figs. 2d and S5). This suggests that, during spontaneous behavior, hippocampus and visual cortex may act as functional ‘hubs’ (Fig. 2e).

While many subspace dimensions were coupled with multiple regions, other dimensions were more exclusive. For example, somatosensory neurons were most predictive of activity in the first subspace dimension of whisker cortex (Fig. S6a). In general, higher subspace dimensions integrated activity from fewer regions. Overall, 71% of subspace dimensions 3 and 4 and 97% of dimensions 9 and 10 were predominantly influenced by one other recorded region (CI: 56%-81% and 88%-100%, respectively; Fig. S6a). These results did not depend on the exact model used; similar results were seen when comparing subspaces across independent models (Fig. S7; see methods).

Different degrees of integration for different dimensions may allow the brain to control how broadly information is shared. Representations along early dimensions will tend to be shared broadly, while higher dimensions will be more specific. Indeed, early subspace dimensions were most strongly correlated with motor activity (Fig. 2f,g), consistent with the idea that motor information is shared widely^6,11^.

## Subspace networks are multiplexed across the cortex

So far, our results suggest brain regions are functionally connected through a shared subspace and that each dimension of this shared subspace is connected with a network of brain regions. However, even with multiple Neuropixels probes, we have a limited view of the network of brain regions that are connected to an individual subspace dimension. Therefore, in order to visualize the broader network, we combined widefield imaging and electrophysiological recordings.

Widefield calcium imaging captured the activity of populations of pyramidal neurons in the superficial layers of dorsal cortical regions^25^. In order to match timescales across imaging and electrophysiology, we estimated the neural activity underlying the calcium signal (Figs. S8-S10; see methods). Then, we correlated the moment-by-moment fluctuations in neural activity along each subspace dimension of a brain region, as measured by electrophysiology, with fluctuations in neural activity across the entire cortex, as measured by the widefield imaging (Fig. 3a; after removing evoked responses, see methods). The resulting maps visualize the network of cortical regions that co-varied with neural activity along each subspace dimension of a region (Fig. 3b; additional examples in Fig. S11; all maps were thresholded at significance, p<0.05, corrected for false-discovery rate, see methods). These maps reflect the ‘subspace network’ of cortical regions that are functionally connected to a dimension of a region’s shared subspace.

**Figure 3.**
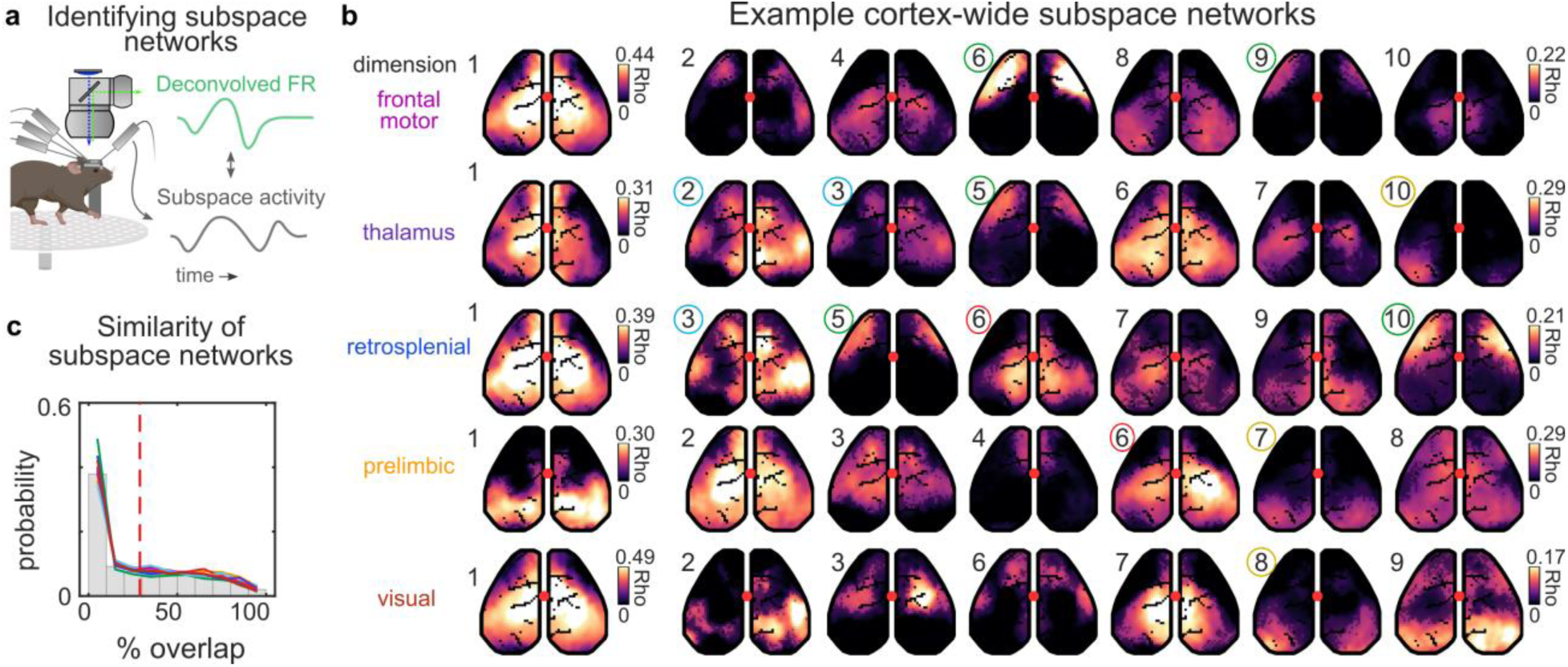
Subspaces engage independent but overlapping cortical networks. **(a)** Electrophysiology was combined with simultaneous widefield calcium imaging of population-level neural activity across cortex. Variance in neural activity projected along each subspace dimension was correlated with imaging signal to identify the subspace-associated cortical networks. To better match the time constants of imaging and electrophysiology, we used a feedforward neural network to estimate the neural activity underlying the imaged calcium signal (see methods). **(b)** Representative cortical maps for different dimensions of subspaces from an example dataset. Rows are different regions. Numbers indicate the subspace dimension corresponding to each map. Colored circles around numbers indicate cluster identity of subspace networks. Color intensity of map indicates the strength of correlation between fluorescence activity at that pixel and the spiking activity along a given subspace (see methods). Transparency (alpha) of color is thresholded at significance such that non-significant pixels are more transparent. For visualization, color is scaled independently for first network versus subsequent networks. **(c)** Similarity of cortical networks. Similarity (x-axis) was computed as the percentage of overlap in significant pixels between cortical maps across all subspace dimensions for a region (see methods). Y-axis shows the fraction of all compared subspaces. Gray bars show all data (across regions, n=630 dataset). Colored lines show mean distribution of each brain region (n=8) across datasets.

There was diversity in the structure of subspace networks. Consistent with our electrophysiology results, the first subspace dimension of most regions was correlated with a broad network that spanned the majority of the dorsal cortex (Fig. 3b, first column; 75.4% of pixels, CI: 73.3-77.2%). In contrast, subsequent subspace dimensions engaged more localized networks of cortical areas (Figs. 3b and S12). Some of these subspace networks followed known functional boundaries, such as anterior somatomotor networks^26,27^ (Fig. 3b, green circles), while others followed anatomical boundaries, such as primary visual cortex (Fig. 3b, yellow circles). Yet, other subspace networks were novel, bridging traditional borders (Fig. 3b, red circles).

Many subspace networks had similar topographies (Fig. 3b, highlighted by colored circles). For example, the subspace network associated with dimensions 6 and 9 in frontal motor was similar to dimension 5 in thalamus and dimensions 5 and 10 in retrosplenial (Fig. 3b, green circles; dimensions 6 and 9 of frontal motor shared 67.5% of pixels, p<0.001 above chance, one-sided binomial test, see methods). Likewise, the subspace network of dimensions 2 and 3 in thalamus was similar to dimension 3 in retrosplenial (Fig. 3b, blue circles; shared 86.7% of pixels, p<0.001). Overall, clustering found 13 categories of subspace networks (categories reflected in color circles in Fig. 3b, see Fig. S13a-e for clusters; clustered across all recordings and regions, see methods).

When two (or more) dimensions within the same region had the same subspace network, it suggests these subspace networks were multi-dimensional (e.g., dimensions 6 and 9 in frontal motor cortex). Overall, we found 38% of subspace networks were multi-dimensional within a region (Fig. S13f; more than expected by chance, p=2e-5, permutation test, see methods). Multi-dimensional subspaces could increase the information capacity of the subspace network, allowing more complex representations to be communicated.

While some subspace networks had similar topographies, many were distinct from one another. For example, in frontal motor, the subspace networks for dimensions 4 and 8 were significantly non-overlapping with dimensions 6 and 9 (average overlap between pairs = 2.3%, all p<0.001, below chance, one-sided binomial test). Overall, any two subspace networks overlapped on an average of 28.9% of their significant pixels. However, there was a wide distribution of overlap across different pairs of subspace networks (STD=28.9%) and a considerable fraction (37.9%) of pairs overlapped in less than 10% of their pixels (defined as <10% overlap; Fig. 3c). The diversity of subspace networks is also reflected in the diversity of clusters, many of which were restricted to a few regions and had little overlap with other subspace networks (Fig. S13b).

Each individual subspace network tended to involve a few distinct regions (Figs. 3b and S13b), but the set of all subspace networks evenly tiled the cortical surface (Fig. 4a). On average, each cortical area was significantly involved in three of the subspace networks from each target region. The partially-overlapping, yet distinct, nature of the subspace networks reflects their multiplexed nature. To visualize this, Figure 4b shows the ‘barcode’ of subspace networks that involves each cortical area for a given target region. For example, in frontal motor cortex, dimension 3 is shared broadly, including both anterior and posterior regions (blue triangles, Fig. 4b, top left), while dimension 4 is restricted to posterior regions (green triangles) and dimensions 2, 5, and 6 are restricted to anterior regions (red, purple, and orange triangles). In this way, different dimensions of the neural representation within frontal motor cortex can engage different networks of cortical areas. From a different perspective, this means that anterior and posterior cortex will have different ‘views’ of the representation in frontal motor cortex: some of the representation will be shared (i.e., what is represented along dimension 3) while other parts of the representation will be distinct to each region (i.e., what is represented along dimensions 2, 4, 5, and 6). Similar patterns were observed in other regions (Fig. 4b). Altogether, these results suggest subspace networks may allow communication between regions to be multiplexed, as different dimensions of neural activity within a brain region are coupled to distinct, yet overlapping, networks of brain regions.

**Figure 4.**
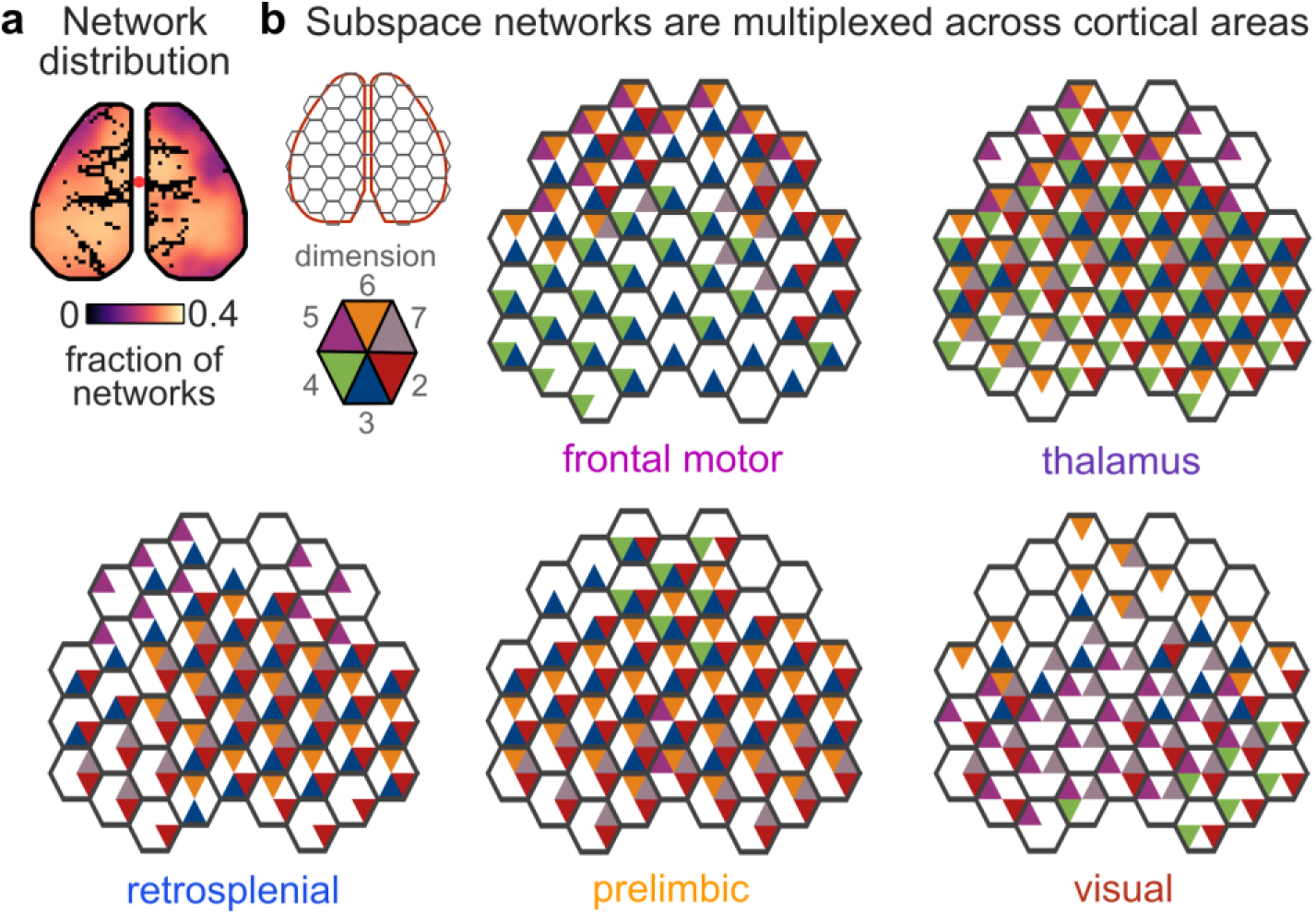
Cortex-wide subspace networks are multiplexed. **(a)** Subspace networks uniformly involved regions across cortex. Map shows the average fraction of subspace networks that were significantly correlated with each pixel (out of 10 possible maps per subspace, averaged across datasets from all target regions, n=630). **(b)** Honeycomb plots showing multiplexing of subspace networks across cortical areas. Each hexagon shows the dimensions of a subspace that significantly engaged over 25% of pixels within that hexagonal parcel of the cortical surface. For visualization, only a subset of dimensions (2-7) is shown.

## Flow of activity between brain areas is associated with alignment of local activity to subspace networks

Finally, we wanted to test the hypothesis that engaging different subspace networks could influence how neural activity propagated between cortical areas. As expected^3,4,11^, cortex-wide imaging found neural activity was dynamic; at different moments in time, different networks of brain regions were engaged (Fig. S1). Using a convolutional factorization approach^28^, we identified 14 ‘motifs’ of cortex-wide neural dynamics^4^. Each motif reflected a unique spatiotemporal pattern of neural activity that captured how neural activity was flowing between brain regions at that moment in time (likely reflecting different cognitive processes^4,29^; see Fig. S1 and methods). Altogether, the 14 motifs captured 65.8% of the variance in cortex-wide neural activity in withheld data (CI: 65.0-66.6%). Importantly, which motif was expressed changed on a moment-by-moment basis^4,5^, reflecting flexibility in how neural activity propagated between brain regions during spontaneous behaviors.

Although motifs were measured using widefield imaging data, they reflected the underlying engagement of individual neurons (Fig. S1). For example, during motifs A and B, neural activity was increased above baseline in whisker, somatosensory, and retrosplenial cortex (all p<0.001, paired t-test; Fig. 5a, insets). However, the magnitude of neural response differed between the two motifs: somatosensory and whisker cortices were more active during motif A, while retrosplenial cortex was more active during motif B (both p<0.05, paired t-tests). This suggests the two motifs evoked different responses across the three regions. Indeed, as seen in Figure 5b, the co-evolution of neural activity across all three regions differed between motifs. Motif A evoked neural activity in whisker and somatosensory cortex (Fig. 5b, red), while motif B largely evoked activity in whisker and retrosplenial cortex (Fig. 5b, blue). Reflecting the different dynamics, the evoked responses for motif A and B were captured by two independent, nearly-orthogonal, planes within the three-region neural space (Fig. 5b, inset).

**Figure 5.**
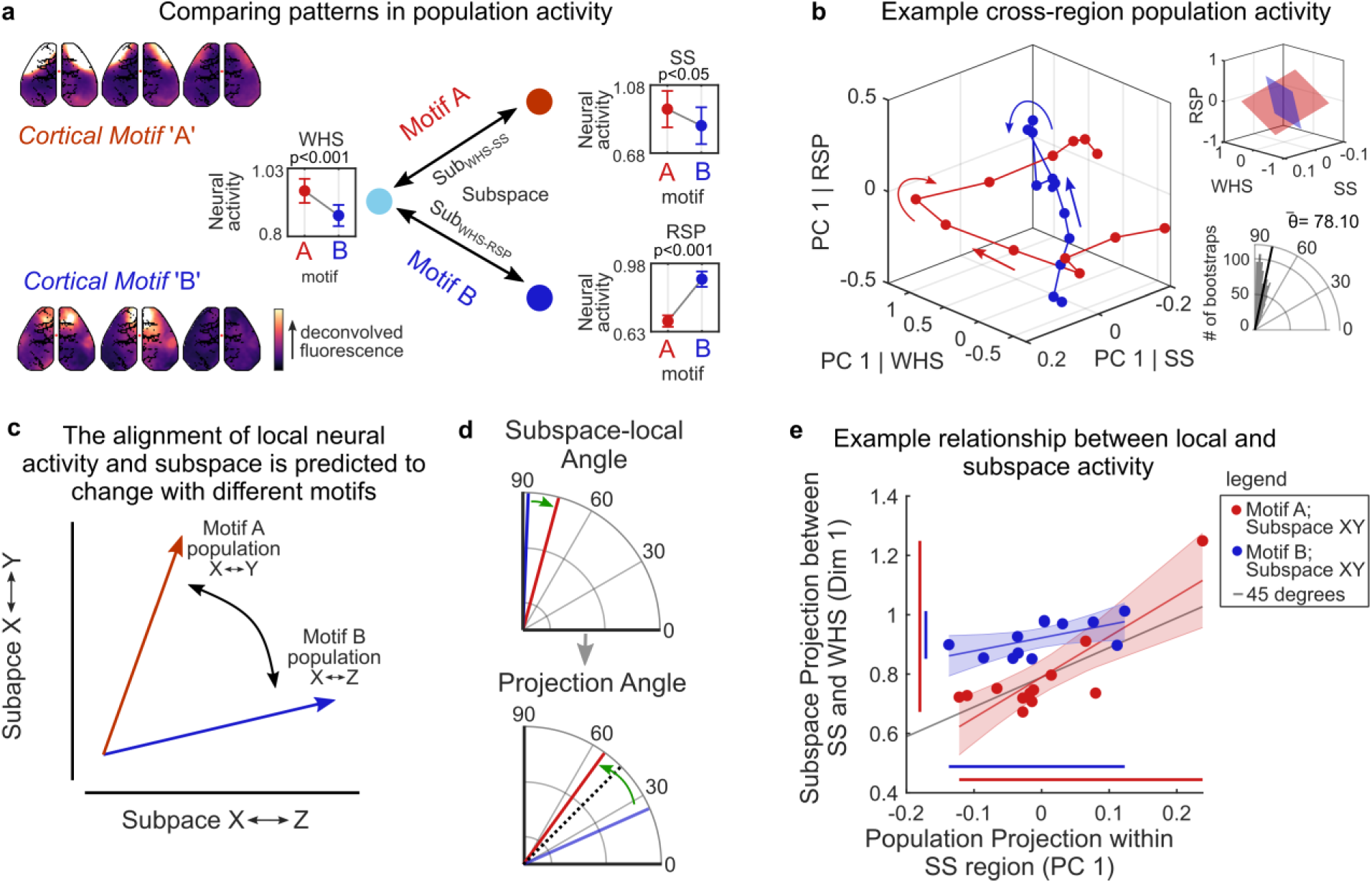
Neural responses are aligned to the shared subspace between two regions when those regions are engaged. **(a)** Left plots show pattern of cortical fluorescent activity during motif A (top) and motif B (bottom). Right plots show the neural response in whisker (WHS), somatosensory (SS), and retrosplenial (RSP) cortex (from electrophysiological recordings) during each motif. Dot and error bars indicate mean and standard error of baseline-normalized spiking activity across neurons during each motif for an example recording (SS, n=29; WHS, n=230; RSP, n=190 neurons, see methods). **(b)** Average projection of neural activity in three-region neural space during motif A (red) and B (blue). Neural activity was projected along the first principal component (PC) of the evoked neural responses in each region (see methods). Arrows show evolution of neural activity over time following onset of each motif (motif A, n=336 occurrences; B, n=266; 133ms timesteps). Top inset shows planes fit to this PC space projection. Bottom inset shows bootstrapped distribution of angle between projection of activity (n=1000 bootstraps across motif occurrences). **(c)** Schematic of proposed mechanism wherein aligning an evoked response to a subspace dimension propagates activity along the associated subspace network. **(d)** Small rotations in the neural representation can lead to large changes in the projection of the neural response along the subspace. Top plot shows a relatively small change in the alignment between the local representation of activity in somatosensory cortex (taken as the first PC) and the SS-WHS subspace dimension when comparing motif B (blue) to motif A (red; taken from example dataset). Bottom plot shows this translates into a large change in the evoked response along the SS-WHS subspace. The efficacy of the evoked response is measured as the angle between the projection along the SS-WHS subspace and the first PC of the local response within somatosensory cortex. Dotted line indicates 45 degrees (i.e., perfect alignment). **(e)** Projection of neural activity during both motifs along the first principal component of activity in SS (x-axis) and the first SS-WHS subspace dimension (y-axis). Greater alignment between the two projections was associated with a larger evoked response; the domain and range of which is shown by red and blue lines along axes. Points show timepoints of evoked response during each motif (as in b). Line and shaded region show least squares fit and 95% confidence bounds, respectively.

Here, we wanted to test the hypothesis that the different neural responses observed during motif A and B reflected the engagement of different subspace networks. In particular, whether aligning the neural response with a specific dimension could engage the associated subspace network (Fig. 5c). This hypothesis predicts the evoked response in somatosensory cortex should be aligned with the somatosensory-whisker (SS-WHS) subspace during motif A, when there is a strong response in somatosensory and whisker cortex, and less aligned during motif B, when there is a relatively weaker response (Fig. 5d, upper). Indeed, this is what we found. During motif A, the angle between the evoked response in SS and the SS-WHS subspace dimension was 73.2°. During motif B, the angle was 85.8° (a significantly greater angle than in motif A, p=0.03, bootstrap, see Methods). In other words, during motif B, the evoked response in somatosensory cortex was nearly orthogonal to the SS-WHS subspace. So, projecting the SS evoked response onto the SS-WHS subspace yielded a relatively small response (Fig. 5e, blue, lines along axes show the extent of the evoked response within somatosensory cortex and projected along the SS-WHS subspace). In contrast, during motif A, the evoked response in somatosensory cortex was more aligned with the SS-WHS subspace, leading to a larger projection onto the SS-WHS subspace (Fig. 5e, red). In this way, a small change in the angle between the neural representation and a subspace dimension (∼12 degrees) can lead to a large change in the projected response (Fig. 5d, lower).

The alignment of evoked responses and subspaces was seen for all pairs of somatosensory, whisker, and retrosplenial cortex during motif A and B. The evoked responses in somatosensory and whisker cortex were more aligned with the SS-WHS subspace during motif A than motif B (Fig. 6a, left two plots). In contrast, the evoked responses in retrosplenial and whisker cortex were more aligned with the RSP-WHS subspace during motif B (Fig. 6a, two right plots). Similar results were observed across all recordings for this trio of regions and pair of motifs (Fig. 6b; p<0.001, bootstrap test). These results show aligning the evoked response to either the SS-WHS or RSP-WHS subspaces is associated with changes in the flow of neural activity between regions (captured by motifs A and B, respectively).

**Figure 6.**
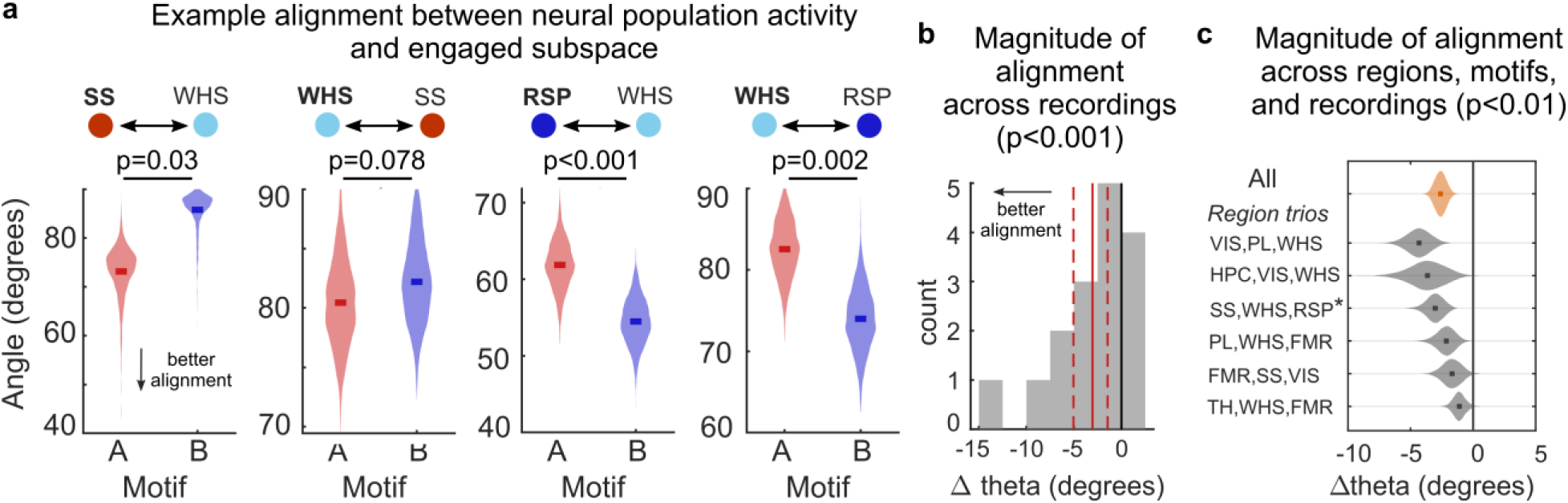
Neural responses were aligned to shared subspaces across regions, motifs, and recordings. **(a)** Bootstrapped distribution of the average alignment between the evoked neural response in either SS, WHS, or RSP (denoted in bold) during motif A or B and the subspace between that region and another region (see methods). Lower angles (y-axis) reflect stronger alignment. n=1000 paired bootstraps. Panels a-f reflect data from the same example recording. **(b)** Distribution of average difference in alignment angle between motif A and B across recordings for SS, WHS, and RSP (n=16 comparisons). Difference in angle was computed relative to the motif with greater engagement of a target brain region, i.e., motif A minus B for comparisons involving SS, but motif B minus A for comparisons involving RSP. Thus, negative values indicate that the motif with greater engagement exhibited better alignment between local neural activity and subspace activity. Vertical red lines show mean (solid) and 95% CI of mean (dashed). **(c)** Bootstrapped distribution of difference in alignment from multiple pairs of motifs and trios of brain regions across recordings (see Fig. S14). Orange violin shows the overall alignment across all comparisons (240 alignment comparisons, see Methods). Overall, the local neural representation and the associated subspace dimension were more aligned when the stronger motif was engaged (p<0.001). Gray violins show alignment distribution for example pairs of motifs/trios of regions (n=32-48 comparisons each, see methods). Individually, neural activity during all compared motifs/regions exhibited significant alignment to the associated subspace. Asterix indicates the example motif pair/region trio used in panels a, b and Figure 5. p-values estimated with bootstrap test relative to zero change in alignment, n=1000 bootstraps.

This was a general phenomenon. A similar pattern of results was seen for six different sets of triads of regions and pairs of motifs (Fig. 6c and S14; p<0.001, bootstrap test; region triads collectively included all 8 recorded regions, see methods). Control analyses confirmed this was not due to changes in the subspace networks. The subspace networks were stable across motifs: independent regression models fit during one motif generalized well to other motifs, explaining 45.2% of the variance in neural activity within the shared subspace (Fig. S15a-b; CI: 44.8-45.5%; see methods). Similarly, the cortex-wide maps of subspace networks were similar between motifs (Figs. S13d and S15c; average of 63.48% overlap between motifs, CI: 63.40-63.56%). Finally, alignment was observed regardless of whether subspaces were defined using neural activity during either motif (see methods).

Our results suggest aligning neural responses with different subspace dimensions may provide a mechanism for controlling how information flows between brain regions. Previous work has shown neural representations within a region can dynamically transform from one subspace to another^30,31^. These transformations depend on the task, suggesting they are under cognitive control^32^. Therefore, transforming representations to be aligned with different dimensions of the shared subspace could engage different subspace networks, and flexibly change how the encoded information is routed to other brain regions (Fig. 5c).

Importantly, because subspace networks are multiplexed, the control of information flow does not have to be all-or-none. Classic models suggest cognitive control acts by propagating all of the information represented in one region to all coupled brain regions. Subspace networks could allow for more nuanced control. Brain regions can represent multiple pieces of information simultaneously, encoding each in a different dimension within the local neural space^31–33^. When these dimensions are aligned with different dimensions of the shared subspace, then the represented information could propagate to different subspace networks. For example, a coarse action signal represented in frontal motor cortex could be broadcast widely by aligning it to the first subspace network while, at the same time, a detailed, efferent copy, of the motor representation could be shared with somatosensory regions by aligning it to subspace networks six and nine (Fig. 3b). In this way, the multiplexed nature of subspace networks may allow different components of a representation to be routed to different regions.

## Acknowledgments

We thank the Buschman lab for their feedback throughout this project and the Princeton Laboratory Animal Resources staff for their support. This work was funded by a grant from SFARI 670183 (T.J.B.), NIH NJCATS Award TL1TR003019 (C.J.M), and R01MH126022 (T.J.B).

## Author Contributions

Conceptualization: TJB, CJM, AL, CIJ, ST; Experimental Design: TJB, CJM; Surgeries and Data Collection: CJM; Analysis and Visualization: CJM, TJB; Writing: CJM and TJB; Editing: TJB, CJM, AL, CIJ, ST; Funding Acquisition: TJB, C.J.M; Supervision: TJB

## Methods

All experiments and procedures were approved by the Animal Care and Use Committee (IACUC) of Princeton University and were carried out in accordance with the standards of the National Institutes of Health.

## Data and Code Availability

Upon acceptance, pre-processed, deconvolved, widefield images, and spike-sorted neural data will be made publicly available through an online data repository and code for figure generation will be publicly available on our GitHub repository (https://github.com/buschman-lab).

## Mice

Experiments were performed on three adult (>8 weeks old) male (N=2) and female (N=1) mice expressing GCaMP6f in cortical excitatory neurons (Thy1-GCaMP6f line^34^). Each mouse was recorded twice for a total of n=6 recordings. Previous studies in animals expressing GCaMP under the Thy1 promoter have not shown epileptiform activity^5,35^. Consistent with this, we screened all recordings for potential epileptiform activity and no events were observed.

## Surgical procedures

Surgical procedures were performed in two stages. First, the skull was made optically transparent and a headplate was installed to allow for awake, *in vivo* widefield imaging. Second, 6-10 days later, four craniotomies were made to allow for acute electrophysiology.

Headplate implantation and preparation of dorsal cortex for widefield imaging followed previous work^4,5^. In brief, mice were anesthetized and given analgesics (Buprenorphine, 0.1mg/kg; Meloxicam, 1mg/kg) and sterile saline. The skin over the dorsal cranium was shaved, disinfected (betadine and 70% isopropanol), and resected. To make the skull optically accessible, the periosteum was removed, and a thin layer of clear dental acrylic was applied to the skull surface (C&B Metabond Quick Cement System). After drying, the acrylic was polished with a rubber rotary tool tip (Shofu, part #0321; Dremel, Series 7700) and coated with clear nail polish (Electron Microscopy Sciences, part #72180). To allow for head fixation, a titanium headplate with 11mm trapezoidal window was cemented to the skull. Mice recovered, single-housed, in a clean home cage with post-op analgesia (Meloxicam, 1mg/kg 24 hours post-surgery).

In the second surgery, mice were anesthetized and given analgesics (Buprenorphine, 0.1mg/kg; Meloxicam, 1mg/kg; Dexamethasone, 3mg/kg) and sterile saline, and ∼1 mm diameter craniotomies were drilled through the acrylic and bone of the dorsal cortex. No visible bleeding was observed during craniotomies. Exposed tissue was kept submerged in sterile saline until coated with Duragel (Dowsil, part #4680) and sealed with Kwiksil (World Precision Instruments). Mice recovered for multiple days prior to recording, during which they were single-housed in a clean home cage with post-op analgesia (Meloxicam, 1mg/kg or Buprenorphine, 0.1mg/kg; 48 hours post-surgery).

To determine craniotomy location, we used a combination of custom code and brainrender^36^ to model electrode insertion paths. This allowed us to design insertion trajectories that targeted our desired brain areas while 1) maintaining insertion angles that would not obscure the imaging field of view and 2) fitting within the confined space of a single hemisphere. We then cross-referenced these potential trajectories with the location of large superficial blood vessels in each mouse - visualized by a brief imaging session prior to the second surgery - to find four craniotomy sites that were consistently situated above frontal motor (FMR), visual (VIS), somatosensory (SS), and retrosplenial (RSP) regions. These sites were centered approximately [-2mm, 1mm], [3mm, 2mm], [2.6um, 0um], [1.2um, 0.9um] along the anterior-posterior, medial-lateral axis relative to bregma, respectively (positive values reflect posterior and lateral to bregma). Insertion angles for each electrode were approximately [32°, 54°, 42°, and 52°] degrees, respectively. This allowed us to target multiple regions with each electrode: 1) FMR and PL; 2) VIS, HPC, and TH; 3) SS and WHS; and 4) RSP.

## Habituation and Behavior

Animals were habituated to head fixation and running on a horizontal treadmill under the microscope between the first and second surgeries. Over four days the animals were acclimated for 30, 60, 120, and 180 minutes.

As we were interested in understanding how the brain flexibly engages different cognitive computations (and the associated networks of regions), the animals were free to perform a variety of different behaviors. Once habituated, animals spontaneously switched behaviors, including periods of running on the treadmill, grooming, whisking, and relative quiescence.

## Widefield Imaging

Widefield imaging was performed using an Optimos CMOS Camera (Photometrics) through back-to-back 50 mm objective lens (Leica, 0.63x and 1x magnification), separated by a 495nm dichroic mirror (Semrock Inc, FF495-Di03-50×70). Excitation light (470nm, 0.4mW/mm^2^) was delivered through the objective lens from an LED (Luxeon, 470nm Rebel LED, part #SP-03-B4) with a 470/22 clean-up bandpass filter (Semrock, FF01-470/22-25). Fluorescence was captured at 30 frames per second (FPS; 33.3ms exposure) using Micro-Manager software (V1.4) at 980×540 resolution (∼34um/pixel) for 90 minutes. Neuropixel recordings and imaging were aligned to the exposure out signal of the CMOS camera (captured on a NIDAQ PXIe-8381 in SpikeGLX; aligned with TPrime v1.6 available from https://billkarsh.github.io/SpikeGLX/#tprime).

Preliminary experiments revealed that the strobed excitation lights often used during widefield imaging for alternating GCaMP6f excitation (blue) and hemodynamic correction (violet or green) produced a large, saturating electrophysiological artifact. Potential mitigating measures, such as excitation intensity ramping^37^ did not sufficiently remove this artifact. As our analyses relied on spiking data, we prioritized eliminating electrical artifacts. To this end, we opted to not strobe LEDs for excitation of GCaMP and hemodynamic correction. Instead, excitation was maintained throughout the recording session. To mitigate hemodynamic effects of large vasculature, we masked non-neural pixels around blood vessels^4,5^. Slow drift in the imaging signal was corrected by normalizing fluorescence to a rolling baseline variance (described below). These steps, combined with subsequent deconvolution (described below), minimized hemodynamic contributions, as evidenced by the fact that our observed strength of correlation between deconvolved fluorescence and ground truth spiking activity matched previous work that used hemodynamic correction^37^. After normalization we spatially binned our imaging signal to 68×68 pixels (∼135um/pixel) for subsequent analysis.

Widefield recordings were registered within mice across days using rigid registration on user-drawn fiducials (n>10 points) of notable vasculature and craniotomy edges. Recordings across mice were registered using rigid registration on user-labeled skull landmarks.

## Estimating the Neural Signal Underlying Widefield Imaging

One-photon widefield imaging is a useful measure of large-scale neural dynamics. However, the precise relationship between the recorded fluorescence signal and the spiking activity of underlying neural populations remains unclear^25,38^. Furthermore, the temporal relationship between fluorescence and spiking is lagged (due to indicator kinetics), which needed to be corrected in order to accurately map between spiking and cortical activity. Finally, differences between the signal recorded from craniotomies (covered by Duragel) and through bone can change the variability of the recorded signal. Together, these effects could bias analyses. To avoid this, we performed an extensive set of supplemental analyses aimed at identifying the normalization procedure and deconvolution method that best estimated the neural population firing activity from a widefield imaging signal (Figs. S8-10).

As shown in Figure S8, we found imaging signals were most comparable across regions when the fluorescence of each pixel was normalized by its rolling standard deviation (i.e., ΔF/σF, where σF is the standard deviation during a 120s rolling window). By definition, this corrected for differences in signal variance across the cranium as well as between exposed tissue (craniotomies) and intact skull. Variance normalization also improved subsequent estimations of firing rate in comparison to alternative approaches of normalizing by dividing by a baseline fluorescence value (i.e., ΔF/F0).

We then estimated relative firing rate by performing non-linear deconvolution on the normalized fluorescence signal. Specifically, on each recording, we trained a shallow feedforward neural network to predict spiking activity at time point t0 using the one second of fluorescence signal on either side of that timepoint (i.e., t-30 to t+30). This method resolved the temporal offset of the neural signal, better generalized to withheld data, and better recapitulated the log-normal statistics of neural spiking than alternative approaches^37–39^ (Figs. S9 and S10). For this analysis alone, spiking data was binned to 30FPS to match the imaging data and normalized to standard deviation across the recording for deconvolution. As expected, widefield imaging predominantly reflected population-level activity of superficial layers of cortex (∼200-600um; Fig. S10g-h).

## Identifying Spatiotemporal Motifs of Cortex-wide Neural Activity

Our goal was to understand how neural activity flows between brain regions. This requires quantifying these dynamics, allowing us to categorize the cortex-wide flow of neural activity at each moment in time. This allows us to 1) estimate the functional connectivity between regions by removing the evoked response (Figs. 1-4) and 2) understand how changes in neural representations influence the flow of neural activity between regions (Figs. 5-6).

To quantify the flow of neural activity across cortex, we used a convolutional factorization approach^4,28^ to identify recurring motifs in our widefield calcium signal. Each motif captured a unique cortex-wide spatiotemporal patterns of neural activity that lasts for ∼700 milliseconds (examples are shown in Fig. S1f). Different motifs have been associated with different cortical processes. Some motifs capture bursts of activity in sensory cortex thought to reflect the processing of sensory stimuli. Other motifs capture bursts of activity in motor regions thought to reflect the preparation and execution of motor movements. Still other motifs capture traveling cortical waves thought to reflect cortex-wide integrative processes^29^. By identifying when each motif occurred, we were able to parse the stream of spontaneous activity into distinct ‘trials’, each with a different pattern of information flow. Importantly, our approach does this in an unbiased, data-driven manner.

Motif discovery followed previous work^5^ and is schematized in Figure S1a-e. We used convolutional non-negative factorization (CNMF) with spatiotemporal penalty terms to identify unique 1-second recurring spatiotemporal sequences in our widefield signal. Motif discovery was performed on deconvolved imaging data binned to 15FPS and then split into one-minute chunks. Alternating chunks were used for motif discovery and withheld for cross-validation. The resulting n=4311 cross-validated motifs were clustered across all animals using a graph-based nearest-neighbor clustering of the spatiotemporal correlation between all pairs of motifs (Phenograph^40^; with k=12 neighbors used for construction of the graph). This process identified 14 clusters of motifs, each with a unique spatiotemporal pattern. A 15^th^ motif was identified and excluded from subsequent analysis as it reflected an obvious artifact of hair entering the dorsal field of view in one animal on one recording. The average motif from each cluster were refit to the full length of all recordings, providing a measure of the activity of each motif at each moment in time. This refitting revealed that the 14 motifs captured the majority (66% CI: 65-67%) of variance in our cortical signal across all across all animals (consistent with our previous work^4,5^).

To determine when a motif was active, we applied a threshold to the motif’s temporal weightings. Thresholding allowed us to capture the moments when cortical dynamics were well described by the motifs and avoid small transients in motif activity that may reflect noise. To determine the optimal threshold, we swept through 50 linearly spaced threshold values for each motif and identified the threshold at which the threshold-triggered pattern in the deconvolved mesoscale activity around the time of the threshold had the strongest spatiotemporal correlation with the original motif. After identifying the best threshold for each motif, we were able to determine the onset times when each motif occurred. On average, each motif occurred 299 times during a recording session (range 226-451). In other words, each motif occurred once every 11-24 seconds.

## Electrophysiological Recordings

Electrophysiological recordings used four Neuropixels^19^ 1.0 probes (phase 3B2), inserted simultaneously (see above for trajectories). Electrodes were inserted under 2x magnification with micromanipulators (Siskiyou, part# MX-1131). Probes were coated with DiI (ThermoFisher Scientific, item #V22885) prior to insertion to allow for post-hoc histological reconstruction. Probes were lowered to desired depths (1.5-5mm, pre-determined from modelling described above) and allowed to settle for >30 minutes before starting recording.

Recordings were 90 minutes in length. Data was acquired using SpikeGLX (v3.0 available from https://billkarsh.github.io/SpikeGLX/), with tip reference mode and high-pass filtered at 300Hz. After collection, data was re-referenced by subtracting the global average across all channels using CatGT (v2), and spike-sorted using Kilosort^41^ (v2.5). Automatically identified units were then manually curated with Phy^42^ into well-isolated single units and ‘multiunits’ that may have aggregated the activity of multiple neurons.

## Histology and Reconstruction of Electrode Placement

Approximately 2 weeks after recording was completed, animals were transcardially perfused and their brains fixed (24-48hrs in 4%PFA) and cryopreserved (30% Sucrose). Brains were coronally sectioned with a cryostat at 60uM thickness. DiI electrode tracks were imaged with a NanoZoomer (Hamamatsu Photonics). Probe trajectories were reconstructed, following previous work^37^ (http://github.com/petersaj/AP_histology; with minor adaptations). Trajectories were cross referenced with the A/P and M/L positioning of insertion sites derived from widefield images, to determine their final positioning. Correlograms of spiking activity showed strong concordance between estimated anatomical boundaries and functional boundaries in spiking activity (Fig. 1d). In one animal, probe insertion location had to be shifted for two electrodes to avoid large blood vessels: resulting in two recordings missing retrosplenial cortex, and one recording missing whisker somatosensory cortex.

Recorded neurons were grouped by anatomical location, as labeled in the Allen Brain Atlas Common Coordinates Framework^43^ (CCF v3). *Prelimbic* (PL; n=1527) included neurons from CCF parent regions Prelimbic (PL), Infralimbic (ILA), Dorsal Anterior Cingulate Area (ACAd). *Frontal Motor* (FMR; n=1257) included neurons from Secondary Motor Area (Mos). *Visual* (VIS; n=716) included neurons from Posteromedial (VISpm), Anterior (VISa), and Anteromedial (VISam) visual areas. *Somatosensory* (SS; n=833) included neurons from nose (SSp-n) mouth (SSp-m) and unassigned (SSp-un) primary somatosensory areas. *Whisker* (WHS; n=805) included neurons from Primary Somatosensory Barrel Field area (SSp-bfd). *Retrosplenial* (RSP; 640) included neurons from Dorsal and Lateral Agranular Retrosplenial areas (RSPd and RSPagl, respectively). *Hippocampus* (HPC; n=353) included neurons from Dentate Gyrus (DG) and Ammon’s horn (CA). *Thalamus* (TH, n=389) included neurons from the Lateral Group (LAT), Medial Group (MED), and Intralaminar nuclei (ILM) of the dorsal Thalamus and Epithalamus (EPI).

## Identifying Functional Connectivity Between Neural Populations

To quantify the functional connectivity between brain regions, we measured the degree to which neural activity in one region could predict the activity in another region. To avoid large evoked potentials in neural activity from obscuring underlying interactions between brain regions^44^, we fit all models to the moment-to-moment variability in spontaneous spiking activity.

Specifically, we split spiking activity into discrete ‘trials’ based on the 14 recurring cortex-wide motifs observed in the widefield calcium signal (see above). We considered a ‘trial’ to be the 1333ms (10 time-bins) following the onset of a motif. To get the change in activity during a trial relative to baseline, we normalized spiking activity to the 400ms baseline period prior to that trial: FR_trial_ = FR_trial_/(FR_baseline_+1), as in previous work^10^. Equivalent results were found when normalizing by subtracting the baseline. Unless otherwise specified, spiking activity was binned with 133ms time bins (7.5FPS) for all analyses. Binning with 66.7 ms bins (15FPS) gave comparable results.

After identifying the trials, we subtracted the mean activity during all trials of a single motif from both the imaging signal and spiking activity (Fig. S1f). The residual activity captured the moment-to-moment variability in the neural population^21^. This allowed us to quantify how two regions were interacting – if two regions interact, then the fluctuations in neural activity in one region should predict the fluctuations in the other region.

For analyses supporting Figures 1-4, we identified interactions by fitting regression models to predict the spiking variability of each ‘target’ brain area from the spiking variability of all other ’source’ brains regions. A total of 630 models were fit; one for each brain region (n=6-8 regions per recording) during each motif (n=14) and during each recording (n=6). To fit each model, we concatenated neural activity across all instances of a motif, resulting in a two-dimensional matrix of neural activity per brain area (*[instances x timepoints] x neurons*). In analyses supporting Figures 5 and 6, models were fit in a pairwise manner across brain areas, with only one region as the source and another region as the target. This resulted in a total of 4592 models.

All models were fit using reduced rank regression^21,45^ (RRR) in MATLAB. Model fitting followed previous work^21^. RRR performs simultaneous regression and dimensionality reduction by identifying a *m*-dimensional set of predictive dimensions in a source neural population that best predicts the trial-to-trial variability in activity of a target neural population according to a linear model:

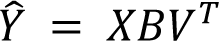

where ^^^𝑌 is a (*N*_trials_ x *N*_timepoints_) by *N*_target neurons_ matrix of predicted activity in the target brain areas. *X* is a (*N*_trials_ x *N*_timepoints_) by *N_source_* _neurons_ matrix of activity in source brain areas (i.e., the independent variable). *B* is a *Nsource* neurons x *m* matrix and *V* is a *N*_target neurons_ x *m* matrix that contain the *m*-dimensional subspace beta weights (β) for the source and target neurons, respectively (i.e., β_𝑖,𝑗_ captures the contribution of neuron 𝑖 to subspace dimension 𝑗). Unless otherwise noted, the first 10 subspace dimensions (*m*=10) were used in all analyses, which captured an average of 90% of the total predictive performance of the fitted models (i.e., “all subspace dimensions” in the main text means the top 10 dimensions).

When fitting β, ridge (L2) regularization was used to avoid overfitting noise in the data. This was particularly important at higher values of *m*, allowing us to better estimate the contributions of higher dimensions. In addition, regularization was important for estimating the total explainable variance captured by the full dimensional model, as shown in Figure 1f (i.e., where *m* = *N*_target neurons_). The strength of L2 regularization was determined through 10-fold cross-validation^21^; the value for regularization was taken as the greatest value for which the mean cross-fold performance (r^2^) was within a standard error of the mean of the peak performance across all regularization values.

Results of model performance (Fig. 1f-h, main text statistics) show cross validated performance of 10 repeats of 10% held-out data. Fitted models were stable across cross-validation folds, i.e., β were highly similar with an average rho between folds of 0.83 CI: 0.82-0.84, for the first 10 subspace dimensions). So, subsequent analyses used models fit to full datasets.

For a given model fit, the positive/negative direction of βs are arbitrary (since the final direction is the product of B and V). So, in order to facilitate the comparison of βs across areas and models (e.g., Figs. 2a-e, S3, and S7), or when projecting activity along β (e.g., Fig. 3b), we used the convention that β weights were oriented such that the majority of neurons had positive weights.

To confirm our results were robust to the analysis approach, we tested two other methods for quantifying interactions. First, as detailed below, we repeated our analyses but reversed the direction of fitting of the reduced-rank-regression models. Instead of predicting the activity of one regions from all other regions, we used the activity in one region to predict activity in all other regions. Second, we repeated our initial analyses using Canonical Correlation Analysis^20,46^ (CCA). CCA is a different, but related, technique for uncovering interregional relationships. CCA found highly similar results to RRR. For example, CCA also found interareal relationships existed in a ‘subspace’ of neural activity, with an average of 5 significant subspace dimensions (significance determined with permutation testing as in previous work^47^).

## Estimating the Dimensionality of Subspaces

To understand the proportion of neural activity that was shared with other regions, we compared the dimensionality of the ‘shared subspace’ within each region to the full dimensionality of population activity within that same region (Fig. 1i-f).

To compute the full dimensionality of the activity within each region, we used a cross-validated measure that identifies dimensions of shared variability within a neural population (shared variance components analysis; SVCA^6^). In brief, neurons within a region were randomly partitioned into two equal groups and their activity was split into two equal sets of timepoints (test/train timepoints). Singular value decomposition of the covariance matrix between the two groups’ activity during the training timepoints yielded a set of covarying dimensions between the neurons in each group. Activity during the test trials was projected along these shared dimensions. The covariance of these projections between the two groups provided a measure of reliable variance for each dimension (i.e., which projections generalize to withheld data). This value was then divided by the average variability within each group to measure the fraction of reliable variance explained by each dimension. The amount of reliable variance of each dimension was then divided by the sum of reliable variance across all dimensions to estimate the percent of explainable variance captured by each local dimension.

Importantly, our finding that the ‘shared subspace’ within each region was significantly smaller than the full space of that region’s population activity was robust to different estimates of dimensionality including both factor analysis^21^ and principal components analysis^47^. Main text results use SVCA because it most conservatively estimated the full dimensionality within a region.

## Using Power-law to Estimate Relative Dimensionality of Full Space and Shared Subspace

Power-law exponents for the decay in dimensionality shown in Figure 1j-k were computed by fitting linear models to the log of the percent of variance explained across dimensions (estimated with SVCA, as described above).

## Estimating the Strength of Interactions Between Brain Regions

The contribution of each neuron to predicting the neural activity of a target region is estimated by the magnitude of its β weight (for schematic, see Fig. 2a). By examining the βs of neurons in different regions, we can quantify the relative strength of interactions between each source region and the target region. For example, Figure 2b shows the distribution of βs of neurons predicting the first dimension of activity of the visual region, sorted by decreasing magnitude. The fraction of neurons within a source region that contributed βs of each magnitude is shown in Figure 2b, right (smoothed with a 50-neuron gaussian kernel). Figure 2c shows the cumulative distribution for each region, averaged over all 84 datasets (see also Fig. S6a). To quantify the contribution of a source region to predicting the activity of a target region, we calculated the area under the curve (AUC) of the source region’s cumulative distribution function. A higher AUC indicates the source region contributed stronger βs, relative to other source regions (see inset of Figs. 2c and S6a for examples). In other words, a region with a higher AUC had stronger interactions with the target region. To estimate how many regions had a relatively stronger influence on the target region, we counted the number of source regions with an AUC greater than 0.5. As detailed in the main text, and shown in Figure S6b, earlier subspace dimensions tended to integrate activity from more regions.

To quantify which regions contributed the most to another region, we averaged the AUCs across all subspace dimensions, and all datasets. This allowed us to generate a graph of the strongest interactions between regions (Figs. 2d-e and S5)

## Subspace Dimensions Are Functionally Connected with Multiple Brain Regions

As detailed in the main text, our results suggest each subspace dimension was functionally connected with a network of regions. One concern might be that this is an artifact of fitting a single model that predicts the activity in one target region from all other source regions (note: regularization discourages this). Therefore, we tested whether similar results were seen when fitting a set of independent models that estimated the ‘projection vectors’ from one source region to each target region (Fig. S7a). If the same source representation was functionally connected to a network of target regions, then the projection vectors for different target regions should be correlated (despite the models being fit independently). Indeed, we found strong correlations in many source regions. For example, the vector of hippocampal activity influencing the first dimension of somatosensory cortex was strongly correlated with the vector influencing the first dimension of prelimbic cortex (Fig. S7b; r=0.65, p<0.001 bootstrap test versus zero correlation). Similarly, the projection vectors from frontal motor to the third dimension of the visual and somatosensory subspaces were significantly correlated (Fig. S7c; r=0.26, p<0.001 bootstrap test). Overall, 30.4% of vectors across all regions were significantly correlated, suggesting that many source regions shared their projections across target regions.

However, it is important to note that not all projection vectors were correlated (Fig. S7d). This reflects the fact that there were multiple dimensions within a source region that influenced different networks of regions. To quantify this, we measured the effective dimensionality of the projection vectors:

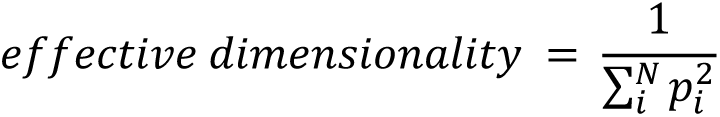

where 𝑝_𝑖_ is the percent of variance explained by each of the *N* principal components of the projection vectors (Fig. S7e). The null hypothesis is that the projection vectors are random and, thus, uncorrelated across regions. Therefore, to generate a null distribution, we randomly permuted the β weights across projection vectors and recalculated the effective dimensionality of each permutation (Fig. S7e). Consistent with partially-shared projections, the observed projection vectors had a dimensionality that was 45.4% of what was expected by random, uncorrelated, projections (Fig. S7e; CI: 44.6-46.2%, p<0.001). Interestingly, although all regions were less than random, there was variability in the dimensionality of the projection space across regions (Fig. S7f). Frontal regions, including frontal motor and prelimbic regions, exhibited the greatest degree of sharing, spanning only 40% of the space (39.2% CI: 38.0-40.4% and 42.2% CI: 40.5-43.9%, respectively). This suggests representations in frontal regions are projected widely. In contrast, hippocampal and retrosplenial regions exhibited the least sharing, spanning 50% of the space (52.3% CI: 50.1-54.6% and 50.3% CI: 47.7-52.8%, respectively; both less than frontal motor and prelimbic at p<0.001, bootstrap test). This suggests HPC and RSP, while having broad impacts on other regions (Fig. 2), tend to have more independent projections to other regions.

## Testing Whether Different Subspace Dimensions Are Functionally Connected with Shared or Independent Populations of Neurons

Our results show there is a subspace of neural activity within each brain region that is functionally connected with other regions (Fig. 1). Different subspace dimensions within a target region are connected with different networks of source regions (Figs. 2-4). Within the target region, each dimension of the subspace was supported by a population of neurons. We were interested in understanding how the network of neurons associated with each subspace dimension related to other subspace dimensions.

Our goal was to discriminate between two hypotheses. First, one might expect that independent sub-populations of neurons support different subspace dimensions (Fig. S3a, upper). Consistent with this, anatomical tracing has found individual neurons send axonal collaterals to a constellation of brain regions, with different sub-populations of neurons projecting to different target regions^43,48^. Similarly, sub-populations of neurons have been found to be correlated with different cortex-wide networks of regions^49^. Different populations of neurons may even carry different types of information; for example, neurons in primary somatosensory cortex (S1) projecting to either secondary somatosensory cortex (S2) or primary motor cortex (M1) are biased to carry different types of somatosensory information^50^.

Alternatively, the same population of neurons could support multiple subspace dimensions (Fig. S3a, lower; either within or between brain regions). Importantly, if the representations associated with each subspace dimension exist along orthogonal dimensions within the local population, then this would still allow different information to be projected to different subspace dimensions. Consistent with this hypothesis, previous has shown neural representation of sensory stimuli^51^, short-term memories^31^, motor movements^11^, and task representations^52^ are distributed across the entire neural population.

To discriminate these hypotheses, we quantified the distribution of β weights associated with pairs of subspace dimensions (i.e., the vectors 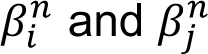 for all neurons 𝑛 in the recorded population 𝑖 𝑗 of 𝑁 neurons for subspace dimensions 𝑖 and 𝑗). The two hypotheses make different predictions on how the β weights should be distributed across neurons. If each subspace dimension was influenced by an independent sub-population of neurons, then we would expect neurons to contribute to one subspace dimension but not the other. In other words, β weights should be large for dimension 𝑖 and small for dimension 𝑗. In a two-dimensional plot of the absolute magnitude of β weights, these neurons would have β weights lie along the x-axis or the y-axis (see Fig. S3b, left, for schematic). Alternatively, if both subspace dimensions rely on the same population of neurons, then the β weights should be evenly distributed across the x-y plane (Fig. S3b, middle). Finally, if the same neurons contribute to both subspace dimensions, then the β weights should cluster around the diagonal (Fig. S3b, right). We used two statistics to quantify the distribution of β weights and, thus, discriminate these hypotheses.

First, we examined the angular distribution of β weights to each pair of subspace dimensions. The angle of the β weights was calculated for each neuron and for each pair of subspaces (taken as 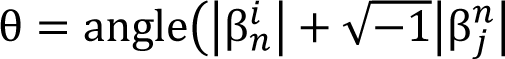), using the *angle* function in MATLAB). To ensure the β weight distribution was not influenced by the number of neurons or the average firing rate of a region, the β weights within each subspace dimension were normalized by their standard deviation. The distribution of angles was measured across all simultaneously recorded neurons (binned into steps of 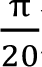). A distribution was estimated for each pair of subspace dimensions, during each motif, 20 and for each recording session. Figure S3c shows the angular histogram for each region; averaged across all subspace dimensions, motifs, and recordings sessions (similar results were seen for individual motifs and recording sessions). Note, all pairs are considered, including mirror opposites (i.e., (𝑖, 𝑗) and (𝑗, 𝑖)) and so this distribution is guaranteed to be symmetrical. This does not affect the above logic.

As seen in Figure S3c, all regions showed a relatively uniform distribution of β weight angles. To estimate the distribution that would be expected given a random distribution of β weights, we randomly shuffled the β weights for each subspace dimension and re-estimated the distribution (Fig. S3c, red lines). In all eight brain regions, the observed distribution of β weights was more clustered along the diagonal than expected by random chance. This was not due to the fact that some pairs of subspace dimensions had correlated β weights; the distribution of β weights for pairs of subspace dimensions that were not significantly correlated (i.e., 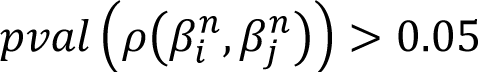, where ρ was Pearson’s linear correlation coefficient) were also uniformly distributed (Fig. S3d, example shown for HPC).

To quantify the distribution of angles, we estimated the curvature of the histogram for each pair of subspace regions. As schematized in Figure S3e, the three models make three different predictions for the curvature of the histogram. If the β weights are from independent subpopulations, then there should be a concentration of β weights along the two axes, resulting in a histogram that is convex. If the β weights are randomly distributed across the population, then we expect a uniform histogram. Finally, if neurons involved in one subspace dimension also tend to be involved in the other subspace dimension, then we expect a concave angle distribution, reflecting the clustering of β weights along the diagonal. The curvature of the histogram was estimated for each pair of subspace dimensions by taking the second derivative of a quadratic equation fit to the histogram: a positive second derivative reflects a convex shape, while a negative second derivative reflects a concave shape (similar results were seen when using the second derivative of the raw data). To test whether the observed curvature was significantly different from chance, we compared the observed curvature to the distribution of curvatures across permuted β weight distributions. This comparison was done for each pair of subspace dimensions, allowing us to calculate a z-score of the observed curvature for each pair of subspace dimensions. Figure S3f shows the distribution of curvatures for each region, across all subspace dimensions. In support of the hypothesis that the same population of neurons influenced both subspace dimensions, the majority of curvatures were below zero for all eight regions (Fig. S3f). A significant percentage of pairs of subspace dimensions were more concave than expected by chance (FMR: 38.3%, PL: 39.0%, WHS: 30.3%, SS: 23.2%, RSP: 21.1%, VIS: 18.6%, HPC:

11.7%, TH: 8.3% of distributions were significantly more concave than chance, p≤0.01 by permutation test; counts in all regions were greater than expected by chance, all p<10^-26^). In contrast, few pairs of subspace dimensions had significantly more convex distributions than expected by chance (FMR: 0.0%, PL: 0.0%, WHS: 0.0%, SS: 0.0%, RSP: 0.0%, VIS: 0.0%, HPC:

0.1%, TH: 0.1% of distributions were significantly more convex than chance, p≤0.01 by permutation test; counts in all regions were less than expected by chance, all p<10^-26^). Similar results were seen when taking only those subspace dimensions with unsignificant correlations in β weights.

Second, we used a receiver operating characteristic (ROC) approach to quantify the distribution of beta-weights across neurons. The β weights were sorted according to their contribution to the first (𝑖) subspace dimension: 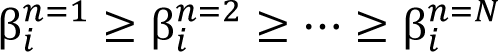 and the ROC was taken as the proportion 𝑖 𝑖 𝑖 of the sorted β weights for the 𝑗 dimension that were above a specific percentile:

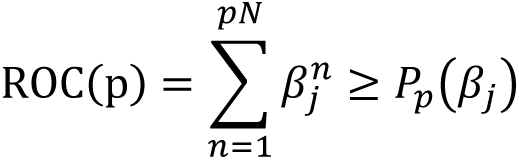

where 𝑃_𝑝_(𝑥) is the value of the p^th^ percentile of distribution 𝑥. To quantify the ROC, we calculated the area under the ROC curve: 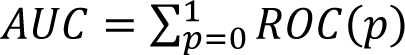. In this way, the AUC quantifies the distribution of β weights. If independent populations of neurons interact with the two subspace dimensions, then the associated AUC will be ¼. In contrast, if the same representation projected to two subspace dimensions (i.e., the β weights are perfectly correlated) then the AUC will be ½. In general, a larger AUC indicates greater overlap in the population of neurons influencing the two subspace dimensions.

The average AUC, across all pairs of subspace dimensions, was ∼0.35 in all eight regions. A majority of pairs within each region had higher AUCs than expected by chance (Fig. S3g), reflecting overlap in the populations of neurons interacting with both subspace dimensions. AUCs greater than chance reflect pairs of subspace dimensions with significantly more overlap in the neural populations than expected by chance (p≤0.01, permutation test). This was the case for a significant number of pairs of subspace dimensions in all eight regions (FMR: 50.3%, PL: 48.6%, WHS: 41.3%, SS: 29.1%, RSP: 29.4%, VIS: 24.9%, HPC: 15.7%, TH: 10.6%; counts in all regions were greater than expected by chance, all p<10^-26^). In contrast, an AUC that is less than chance (p≤0.01, permutation test) would indicate independent populations of neurons. This was the case for significantly fewer pairs of subspace dimensions than expected by chance (FMR: 0.0%, PL: 0.0%, WHS: 0.0%, SS: 0.0%, RSP: 0.0%, VIS: 0.0%, HPC: 0.0%, TH: 0.1% of distributions were significantly more convex than chance; counts in all regions were less than expected by chance, all p<10^-26^)

Interestingly, there was slightly more overlap in the populations of neurons that influenced a pair of subspace dimensions when the two dimensions were in the same target region as supposed to being in two different regions. This was reflected in a significant difference in both the curvature of the β weight histogram and the AUC. The β weight histograms were more concave when comparing two subspace dimensions within the same target region compared to between two different target regions (quantified using a contrast measure (𝑐𝑢𝑟𝑣𝑒_𝑏𝑒𝑡𝑤𝑒𝑒𝑛_ − 𝑐𝑢𝑟𝑣𝑒_𝑤𝑖𝑡ℎ𝑖𝑛_)/ (𝑐𝑢𝑟𝑣𝑒_𝑏𝑒𝑡𝑤𝑒𝑒𝑛_ + 𝑐𝑢𝑟𝑣𝑒_𝑤𝑖𝑡ℎ𝑖𝑛_); FMR: -0.085, PL: -0.091, WHS: -0.062, SS: -0.112, RSP: -0.084, VIS: -0.104, HPC: -0.111, and TH: -0.104). The distributions of curvature were significantly different in all eight regions (FMR: t(174753)=43.64, PL: t(157848)=36.02, WHS: t(155188)=24.47, SS: t(177168)=39.25, RSP: t(135238)=28.70, VIS: t(177168)=37.95, HPC: t(177168)=32.57, and TH: t(177168)=25.86; all p<10^-146^ by t-test). Similarly, the AUCs were significantly higher when subspace dimensions were within the same target region than between regions (contrast: FMR: -0.079, PL: -0.085, WHS: -0.056, SS: -0.098, RSP: -0.075, VIS: -0.094, HPC: -0.098, TH: -0.098; t-test: FMR: t(174753)=-46.68, PL: t(157848)=-36.48; WHS: t(155188)=-25.28, SS: t(177168)=-38.23, RSP: t(135238)=-29.77, VIS: t(177168)=-39.43, HPC: t(177168)=-33.92, TH: t(177168)=-28.66, all p<10^-139^ by t-test).

Altogether, these results support the hypothesis that the same population of neurons is functionally connected with multiple subspace dimensions in other brain regions. We found this was true even when the representations influencing two different subspace dimensions were uncorrelated. Our results are consistent with anatomical studies showing axons from a single neuron can branch, sending collaterals to several brain regions. Furthermore, it is important to note that measures of functional connectivity may include multi-synaptic influences, and so a shared pool of neurons may project to other brain regions by acting locally on different sub-populations of neurons. This would allow the same information to be routed to multiple regions simultaneously, even if the connections between brain regions were anatomically separate.

## Identifying Subspace Networks in Imaging Data

To understand how the neural population within a region was functionally connected with regions across the cortex, we correlated the activity projected along each subspace dimension with the deconvolved widefield fluorescence signal. Activity along a subspace dimension was defined as the dot product of the activity of a region and the β of a subspace dimension (resulting in a *N*_trials_*\*N*_timepoints_ vector). This vector was then correlated with the (*N*_trials_\**N*_timepoints_) *by N_pixels_* matrix of cortical fluorescence activity, yielding a correlation value for each pixel.

To visualize significant cortical networks (Fig. 3b), the observed correlation maps were compared with maps generated from trial shuffled data. We generated an FDR-corrected null distribution of correlation values by taking the strongest correlation across all pixels for each of 1000 trial-permuted maps. Pixels with positive correlations greater than the 95% of this null distribution were taken as significant. The resulting map defined the “subspace networks”.

The similarity of two subspace networks, shown in Figure 3c, was computed as the percentage of pixels that were significant in both subspace network *A* and subspace network *B*:

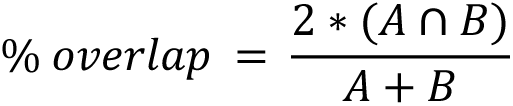

## Clustering Subspace Networks

As shown in Figures 3 and 4, each dimension of neural activity within a brain region was correlated with the activity in a cortex-wide, distributed network of regions. Many of these ‘subspace networks’ were distinct from one another. For example, dimension 1, 2, 4, and 6 of area FMR were associated with different subspace networks (Fig. 3b). However, other dimensions were associated with similar subspace networks (e.g., FMR-6, FMR-9, TH-5, RSP-5, and RSP-10, Fig. 3b). To quantify the distribution of subspace networks, we clustered the subspace networks associated with the first 10 dimensions in each brain region, across all datasets. To minimize the impact of noise, only subspace networks that included significant correlations with more than 5% of the pixels were used. After exclusions, a total of 6,279 subspace networks were clustered. Clustering was done using the Phenograph algorithm^40^. Distance between data points was defined as the inverted correlation of the correlation vectors for area 𝑎_𝑖_, dimension 𝑑_𝑖_ and area 𝑎_𝑗_, dimension 𝑑_𝑗_: dist = 1 − 𝑐𝑜𝑟𝑟 (ρ_𝑎𝑟𝑒𝑎𝑖,𝑑𝑖𝑚𝑖_, ρ_𝑎𝑟𝑒𝑎𝑗,𝑑𝑖𝑚𝑗_), where ρ_𝑎𝑟𝑒𝑎𝑖,𝑑𝑖𝑚𝑖_ is a vectorized form of the map of correlation between the electrophysiologically recorded neural activity in area 𝑎_𝑖_ and dimension 𝑑_𝑖_ and the cortex-wide neural activity as imaged with widefield calcium imaging (i.e., the non-thresholded subspace network). Similar results were seen with other metrics of the distance between subspace networks, including measuring the correlation between the z-transformed correlation maps or measuring the city-block distance between the cortex-wide correlation maps, threshold by significance.

We chose to use the Phenograph algorithm as it does not require defining the number of clusters *a priori*; instead, depending only on the parameter *k*, which defines the number of edges connecting each data point to other similar data points. Phenograph is robust to changes in *k*, with a wide range of values providing good clustering results^40^, although, generally, lower *k* values lead to more clusters and higher *k* values lead to fewer clusters. Our null hypothesis is that each subspace network is unique. Therefore, we chose *k*=16 to be conservative in our estimation of the number of clusters. However, our results did not depend on the value of *k*; similar results were seen across several values of *k*. Furthermore, alternative clustering algorithms, such as identifying the *n* exemplars that minimized distance of each data point to an exemplar, yielded similar results.

After Phenograph clustering, a self-organizing map approach was used to assign individual subspace networks to the most similar cluster. On each iteration, the median subspace network was calculated for each cluster. Each individual subspace network was then assigned to the cluster with which it had the highest correlation to the median. This process was repeated until no subspace networks changed clusters. Combining the Phenograph and self-organizing map algorithms improved the quality of the clustering by increasing the within cluster similarity.

Clustering identified 13 clusters of subspace networks. Figure S13a shows all 6,279 subspace networks projected into a two-dimensional space (using the tSNE algorithm^53^). Each point represents a different subspace network, colored by the identity of its cluster. Each cluster captured a different network of regions – Figure S13b shows the median subspace network correlation map for each cluster. Subspace networks were more similar within a cluster than between clusters (Fig. S13c shows similarity map between all subspace networks).

As noted in the main text, many dimensions, both within and between regions, had similar subspace networks, quantified by the fact that they were members of the same cluster. All regions had subspace dimensions that were correlated with subspace networks belonging to each cluster (Fig. S13d). These results suggest the same subspace network was functionally connected with several brain regions, which could allow a subspace network to integrate information across multiple regions. Note, the distribution of subspace networks across areas was not uniform – the entropy of the distribution was significantly less than expected by chance (p≤0.0002, permutation test). Therefore, while each cluster of subspace networks was engaged by each brain region, they were engaged to different extents by different areas. This is consistent with differences in anatomical connections between regions.

Similarly, the same subspace networks were engaged during different motifs (Fig. S13e). This is consistent with the observation that motifs did not change the underlying structure of interactions, as detailed below.

## Early Subspace Network Dimensions were Correlated with Motor Activity

Recent work has found representations of movements are widely distributed across cortex^6,11^. Motivated by this, we tested whether activity in each subspace network was correlated with motor activity. To this end, we used cameras (PS3 EYE webcam; 640×480 resolution), to track movements of the animal’s nose, whisker-pad, and front shoulder. Regions of interest (ROIs) were manually defined for each region (Fig. 2e). Motion energy was calculated as the average absolute temporal derivative of pixels within all three ROIs (similar results were seen for each ROI independently). Motion energy was then correlated with neural activity projected along each subspace dimension (as detailed above). To account for delays between the signals, the strength of the correlation was taken as the maximum cross-correlation value between the subspace activity vector and behavioral activity vector, up to a 533ms offset between the signals.

The first subspace dimension - and to a lesser extent the second dimension - of most regions was significantly correlated with movements (Fig. 2f). These networks tend to be broad, interacting with many different cortical regions (Figs. 3b and S11). This is consistent with the idea that broad subspace networks may broadcast broadly relevant information, such as motor actions, across cortex.

## Subspace Networks were Stable Across Motifs

As detailed in the main text, one hypothesis is that neural activity engages different regions during different moments in time because the subspace networks themselves change. If true, then we might expect different subspace networks to occur during different motifs. This is not what we found. Instead, we found subspace networks were stable across motifs. To quantify the stability of subspace networks between motifs we tested whether the subspace βs fit on one motif could generalize to another motif. Specifically, we tested the cross-generalization of the reduced rank regression:

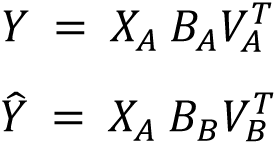

where 𝐵_𝐴,𝐵_ and 𝑉_𝐴,𝐵_ are the βs of the source and target neurons defined during trials of motif A and B. *X*A and Y are the activity of neurons in the source and target regions during motif A, respectively.

For each subspace dimensions, the similarity between motifs was computed as the percent of variance in *Y* explained by ^𝑌^^. However, since subspace dimensions could be re-ordered when fit to different motifs, ^𝑌^^ was computed for each of the first ten dimensions in motif B and reordered to best match each dimension of Y in motif A. This allowed us to test the similarity between motifs irrespective of dimension reordering. The result was a 1xn vector (*S*) of percent similarity between the subspaces for each dimension in motif A. Total percent generalization was then computed by weighting *S* by the amount of explainable variance of the subspace captured by each dimension (1xn vector, *P*):

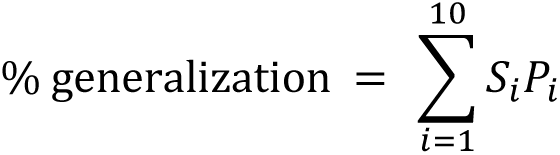

Thus, the total percent generalization reflects the total amount of variance of activity in a subspace that can be explained when generalizing across motifs.

Consistent with the hypothesis that subspaces were stable across time, we found that regression models fit during one motif generalized to capture over 45% of the variance in neural activity of the other motifs across target regions and datasets (Fig. S15c).

## Comparing Evoked Responses in Three-Region Neural Space

As shown in Figures 5a and S1, motifs reflected neural activity in multiple cortical regions. Different motifs were associated with different magnitudes of neural response in each brain region. To visualize the pattern of neural activity in a triad of brain regions, we projected the neural activity in all three regions into a three-dimensional space defined by the first principal component (PC) of each region (Fig. 5b). For these analyses, we were interested in comparing the evoked response across different motifs. For this reason, PCs were fit to the raw neural activity evoked during the motifs (a *N_neurons_ by* [*N*_trials_\**N*_timepoints_] matrix). This is in contrast to the analyses of subspaces, which were defined using RRR on the trial-to-trial variability in firing rate (detailed above).

PCs were fit to the neural activity during one of the two motifs (chosen at random). Similar results were seen when fitting PCs to neural activity from the alternative motif or from both motifs (the latter of which required dropping data to balance occurrences between motifs).

As seen in Figure 5b, neural activity evolved differently during the two motifs. To quantify this, we fit planes to the evoked response within the three-dimensional co-response space. Planes of best it were estimated from the first PC fit to the evoked response across all three regions (Fig. 5b, upper insert). The angle between planes was taken as the angle between the vectors normal to each plane (Fig. 5b, lower insert).

## Calculating Alignment Between Subspace Activity and Local Neural Activity

Finally, we tested the hypothesis that aligning the neural response with a particular subspace dimension would engage the associated network of brain regions (Figs. 5 and 6). If true, then the alignment of the evoked neural response and the subspace should be related to how neural activity flows across cortex. In other words, alignment should change as a function of the motifs.

To test this hypothesis, we measured the angle between 1) the evoked activity in a region and 2) the vector of β weights estimating the subspace between two regions for different motifs. The evoked neural activity within each region was estimated by the first principal component of the neural response during each motif 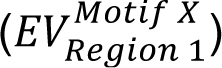. Subspaces were defined using reduced rank regression but fit to each pair of regions independently. This allowed us to isolate the interactions between a pair of regions. As above, models were fit to the trial-to-trial variability in firing rate during each motif. This resulted in a subspace 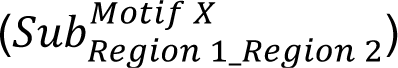 for each region pair during each motif.

For example, the panels in Figure 5a show the following angles (from left to right). The left panel shows the distribution of angles between the WHS-SS subspace, defined using the trial-by-trial variability during motif A 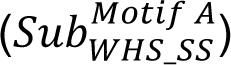, and the evoked response in primary somatosensory regions during motif A (red; 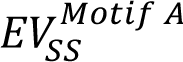) or motif B (blue; 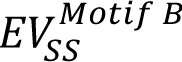). The evoked response during motif A was more aligned with subspace, reflected in the smaller angle. This is consistent with the hypothesis; a stronger neural response was seen in WHS and SS during motif A, which may reflect better alignment of the evoked response and the subspace between WHS and SS.

Similar results were seen for the other pairs of regions in the triad. For example, the second from the left panel shows the angle between the WHS-SS subspace, defined using the trial-by-trial variability during motif A 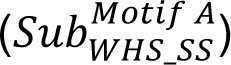, and the evoked response in whisker cortex during motif A (red; 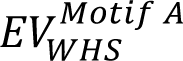) or motif B (blue; 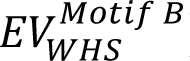). As above, the evoked response and subspace was more aligned during motif A, consistent with a stronger response in the WHS and SS network.

The third from the left panel shows the angle between the WHS-RSP subspace, defined using the trial-by-trial variability during motif B 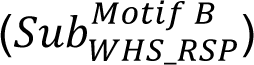, and the evoked response in whisker cortex during motif A (red; 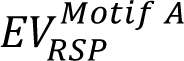) or motif B (blue; 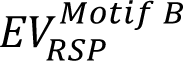). Finally, the rightmost panel shows the angle between the WHS-RSP subspace, defined using the trial-by-trial variability during motif B 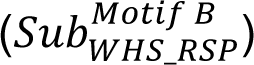, and the evoked response in whisker cortex during motif A (red; 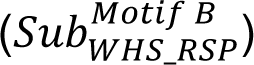) or motif 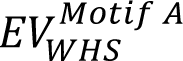 B (blue; 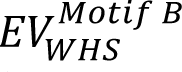). Again, consistent with our hypothesis, the evoked response in both regions were better aligned with the WHS-RSP subspace during motif B, which was the motif that more strongly engaged RSP.

Note that these examples are measuring the alignment of subspaces defined on the trial-to-trial variability of the motif that more strongly engaged a pair of regions (i.e., Motif A for WHS-SS and Motif B for WHS-RSP). Similar results were observed when using the other motif (i.e., Motif A for WHS-RSP and Motif B for WHS-SS). Figure 5c shows the aggregate of alignment angles using both motifs for each pair of regions.

In the main text (Figs. 5 and 6) and supplement (Fig. S14), alignment was calculated for six example triads of regions and pairs of motifs. These were manually chosen based on the pattern of activity seen within the motifs. All pairings were screened to confirm that 1) the motifs engaged all three brain regions (i.e., neural activity was greater than baseline activity) and 2) the magnitude of the evoked response differed between motifs (i.e., each motif preferentially engaged at least one brain region; Figs. 5A and S14). In addition, these example pairings were selected to reflect a distributed sample of the overall data; collectively including all 8 regions and 8 different motifs. Choices of region triads and pairs of motifs were made while blinded to the alignment results and all pre-chosen comparisons were included in calculation of the final statistics (Fig. 6c).

## Statistics and Reproducibility

Unless otherwise noted, we combined across n=14 motifs and n=6 recordings (from three mice) when performing statistical analyses (this follows previous work^21^). Thus, throughout the main text and methods a *dataset* refers to results from all trials of one motif during one recording. When applicable, tests were corrected for multiple comparisons. Permutation tests and bootstrap statistics were performed using 1000 shuffles/resamples. Unless otherwise noted, statistical tests were one-tailed tests of specific predictions.

## Choice of Representative Examples

Our recordings spanned multiple brain regions. To limit bias in data presentation when illustrating general results (e.g., Figs. 1h, 1j, and 2a-c), all examples use data from visual cortex. Similar results were seen in all regions. The choice to use visual cortex was made prior to final analysis.

---

**Supplementary Figure 1:**
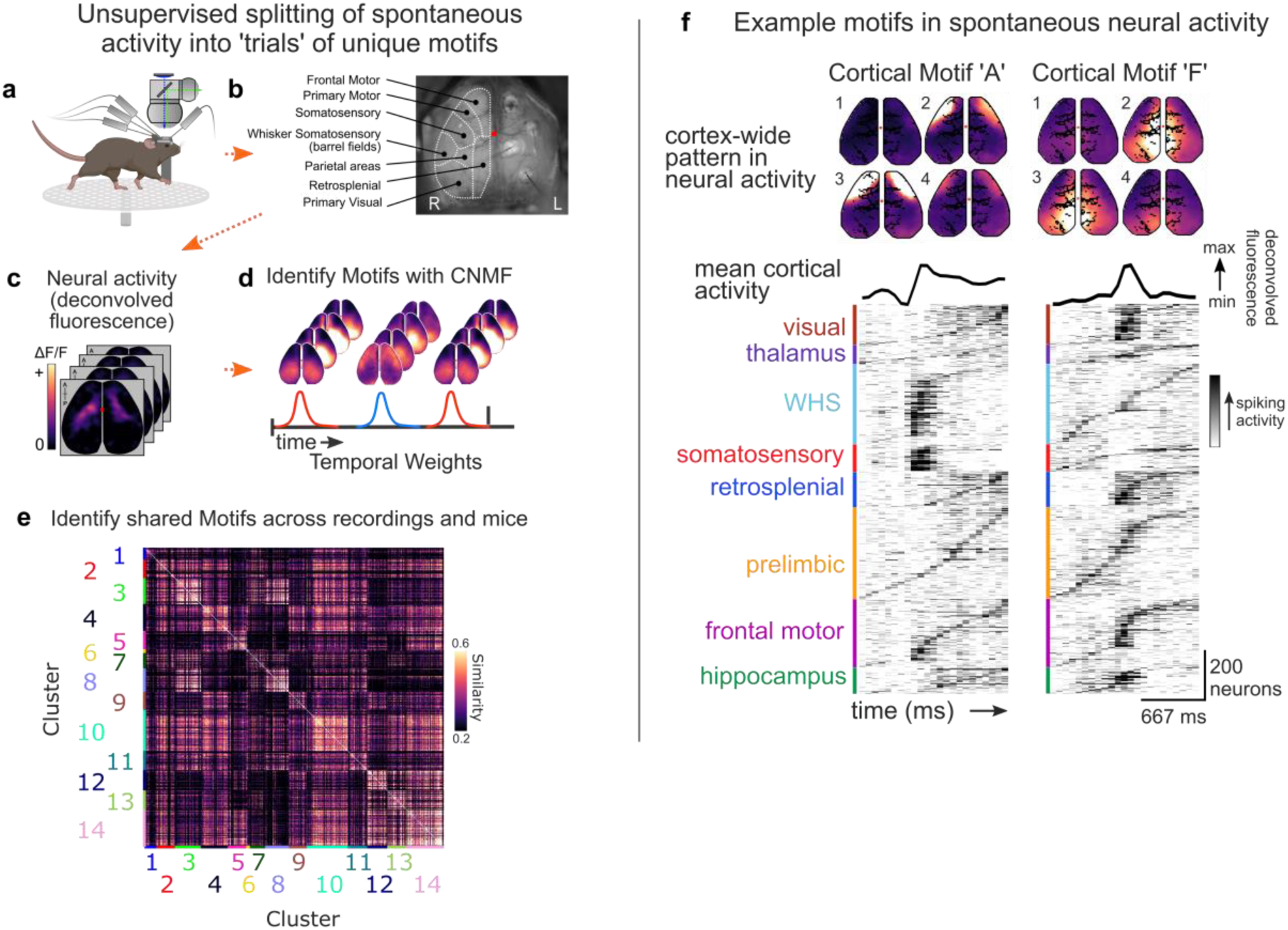
Using motifs to capture repeated spatiotemporal patterns of cortical activity. **(a-d)** As detailed in the main text and methods, we aimed to identify interactions between brain regions by identifying co-variability in their neural activity across a diversity of spontaneous processes. To do so, we used a convolutional factorization approach to split spontaneous neural activity into discrete ‘trials’ based on recurring motifs or cortical activity captured by widefield imaging. **(a)** Schematic of experimental recordings combining multi-region electrophysiology with simultaneous cortex-wide widefield imaging. **(b)** Example field of view of widefield imaging and anatomical parcellation. **(c)** Fluorescence was normalized and deconvolved with a shallow feedforward neural network in order to estimate the underlying neural activity (see methods for details). **(d)** Convolutional non-negative matrix factorization discovered repeated motifs of cortical activity. Each motif captured a unique, ∼1 second, spatiotemporal pattern (top) that can be tiled together to recreate the movie of neural activity (bottom). **(e)** Clustered correlogram showing the spatiotemporal similarity between motifs discovered on individual chunks of recordings. Row and columns correspond to individual motifs. As detailed in the methods, unsupervised clustering revealed 14 shared motifs across animals and recordings. **(f)** Spontaneous activity was split into trial types based on when each of the 14 shared motifs occurred. Cortical maps (top) show the cortex-wide pattern of two example motifs. Black trace shows activity over time of each motif. Panels show the average Peri-Motif Spike Histogram (PMSH) of spiking activity within each brain region. Rows correspond to individual units and are ordered by peak activity for each motif. Therefore, brain areas with a temporally-localized burst of activity mean they exhibited consistent, motif-locked activity across neurons, while areas with more sequential patterns of neural activity are less motif-locked. Color intensity reflect average baseline normalized firing rate across occurrence of motif ‘A’ and motif ‘F’. As noted in the methods, the average PMSH was subtracted from each trial of a motif, yielding the residual moment-to-moment co-variability between neural populations.

**Supplementary Figure 2:**
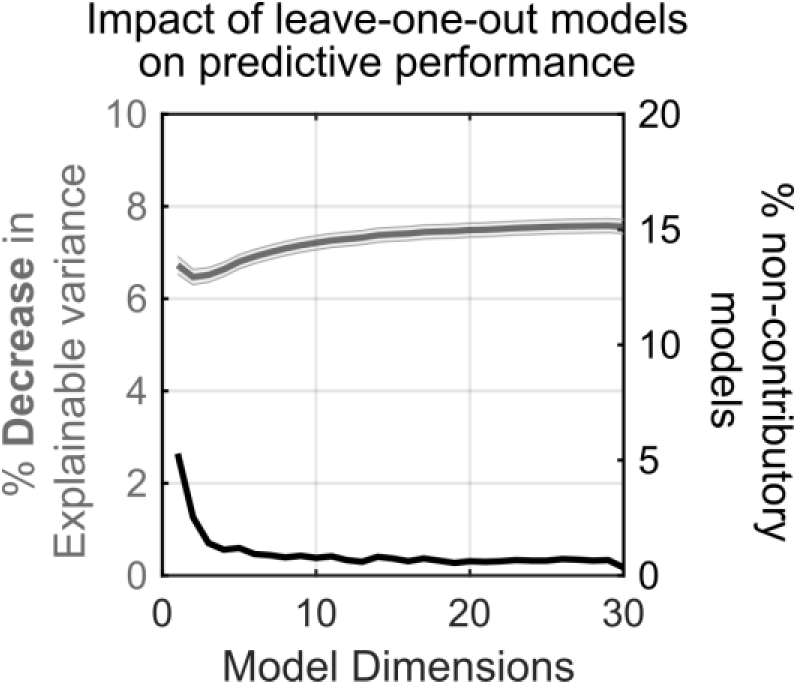
Subspaces integrated activity across a distributed network of brain regions, regardless of the number of dimensions used. We compared the cross-validated performance of reduced rank ridge regression models fit using all brain regions to the cross-validated performance of leave-one-out models fit when excluding any one source region (see methods). Left y-axis and gray line show the magnitude of decrease in performance when excluding a source region. X-axis shows the number of dimensions (rank) of the regression models. Excluding any source region decreased performance between 6 and 8%, regardless of the number of dimensions used (tested up to a rank of 30). Right y-axis and black line show the percentage of models where performance did not decrease when excluding a source region. For all tested dimensions, the vast majority of models (>97%) performed best when using all source regions, consistent with subspaces integrating activity across a distributed network. Lines and shaded regions reflect mean and SE of n=4838 leave-one-out models.

**Supplementary Figure 3.**
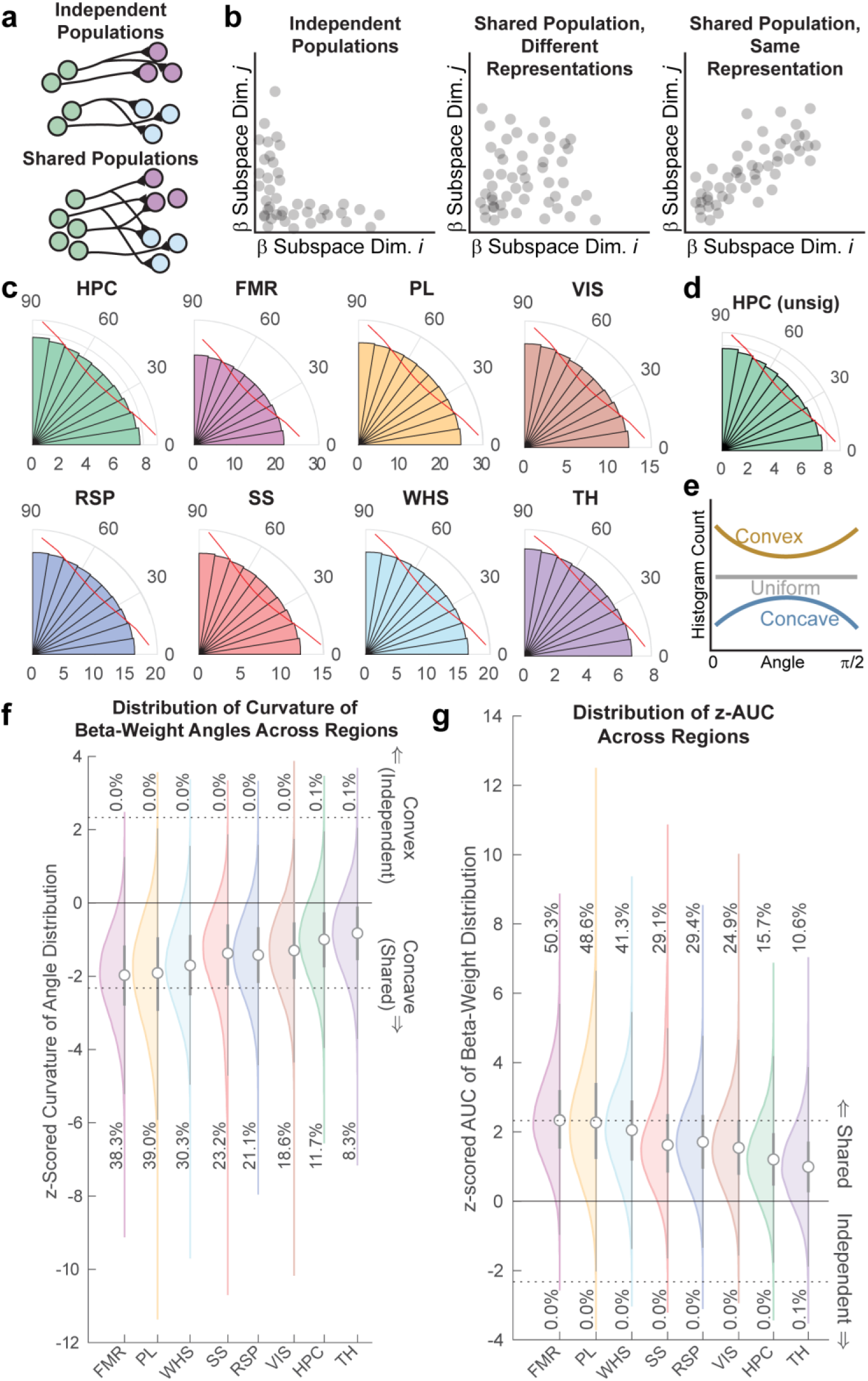
Subspace dimensions relied on overlapping neural populations. **(a)** Schematic of how two different subspace dimensions within a region (green), projecting to two different regions (purple and blue), could either rely on independent populations of neurons (top) or the same population of neurons (bottom). **(b)** These two models make strong predictions about the distribution of the β weights for the two subspace dimensions. If independent (left), then β weights should fall along either of the two axes (i.e., they contribute to subspace dimension *i* or *j*, but not both). If the same population of neurons support both subspace dimensions, this could either reflect independent (orthogonal) β weights for the two subspace dimensions (middle) or the same β weights for both subspace dimensions (right; i.e., they are the same representation). **(c)** Histograms showing the angular distribution of β weights for all eight regions. Red lines show the distribution expected by chance. The distribution of angles is uniform, consistent with the shared, yet independent, model. **(d)** Distribution of β weights for dimensions within HPC that are not significantly correlated. Results are similar to the distribution for all subspace dimensions. Overall, the uniform distribution is consistent with the shared, but independent, model. **(e)** Schematic of the statistic used to measure the shape of the distribution of β weights. Convex distributions (gold) are consistent with independent neural populations, uniform distributions are consistent with shared, but independent, neural populations (grey), and concave distributions (blue) reflect two subspace dimensions engage the same representation. **(f)** Distribution of convex/concave score across all pairs of subspace dimensions and all regions. Very few pairs of subspace dimensions showed convex distributions (independent), suggesting that subspace dimensions relied on a shared pool of neurons. **(g)** Distribution of AUC of β weights across all pairs of subspace dimensions for all brain regions. As detailed in methods, the AUC measures whether the population of neurons supporting each subspace dimension are shared (higher AUC) or independent (lower AUC). Consistent with the histograms of β weights (panel d), almost all pairs of subspace dimensions relied on a shared population of neurons.

**Supplementary Figure 4:**
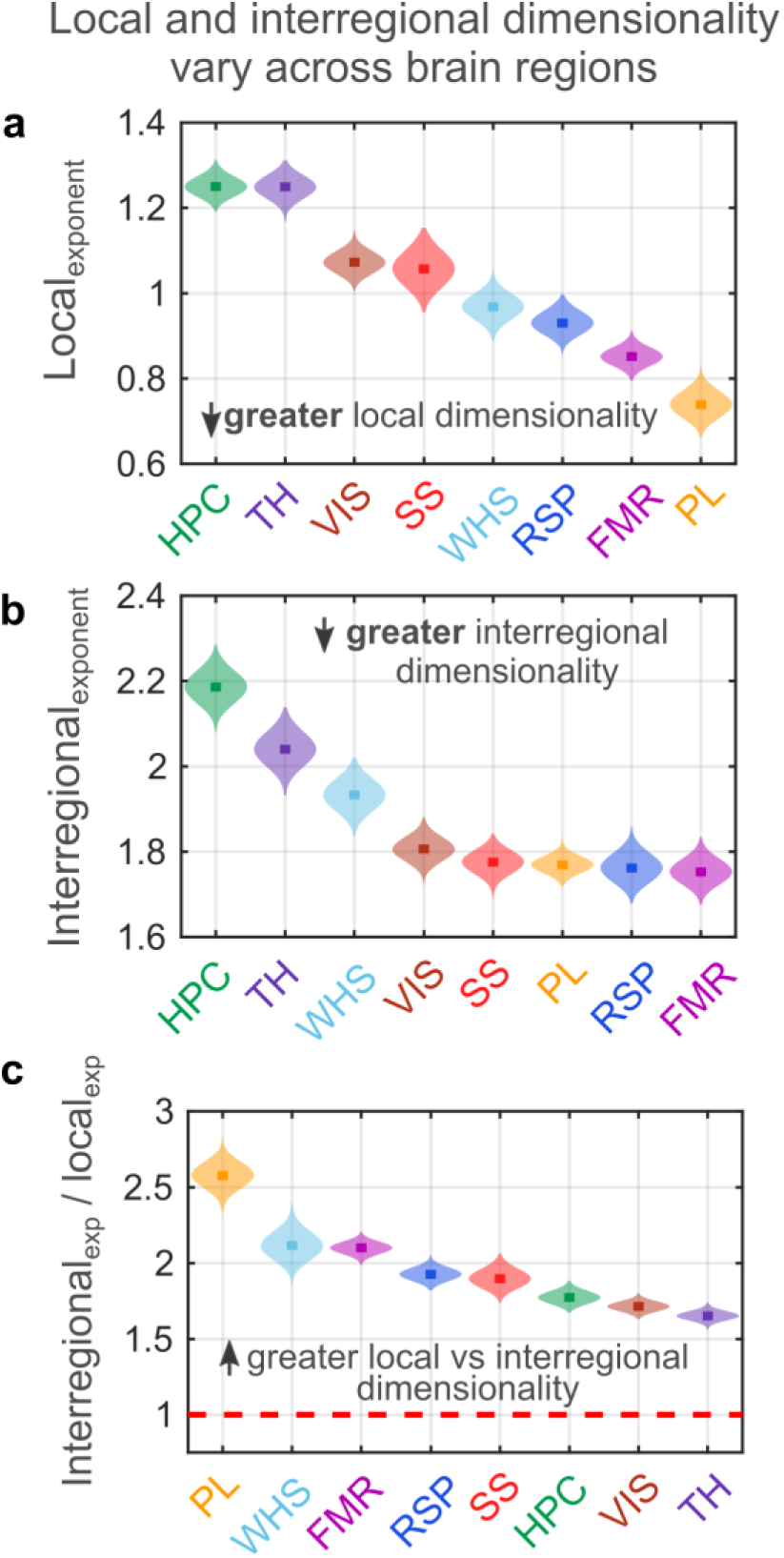
Comparing the dimensionality of local and interregional subspaces. **(a-b)** The variance explained by subsequent dimensions in local or interregional neural activity decreased as a power law (see Fig. 1j). Local and interregional ‘dimensionality’ was estimated as the absolute value of the exponent of these power law fits. A smaller exponent reflects a more gradual decay and thus higher dimensionality. **(a)** Distribution of exponents for fits to local neural activity within each brain region. **(b)** Distribution of exponents for fits to the inter-regional subspace activity between each brain region and all other brain regions. **(c)** Ratio of interregional and local dimensionality (y-axis), shown for all brain regions (x-axis). Greater values along y-axis indicates relatively higher local dimensionality. All distributions reflect n=1000 bootstraps. Colors indicate regions and follow other figures.

**Supplementary Figure 5:**
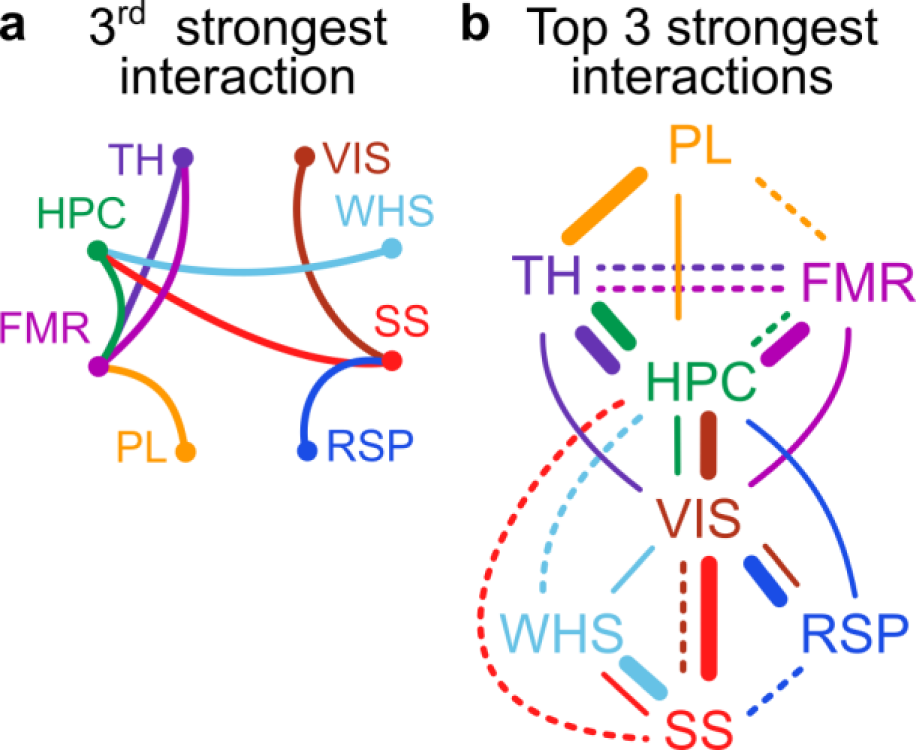
Summary of strength of interactions between brain regions. **(a)** Schematic showing the network of third strongest interactions between areas. Presentation follows Figure 2d. **(b)** Combined network of top three strongest interactions between brain areas. Thick/thin/dotted correspond to the 1^st^, 2^nd^, and 3^rd^ strongest connections, respectively. Presentation follows Figure 2e.

**Supplementary Figure 6:**
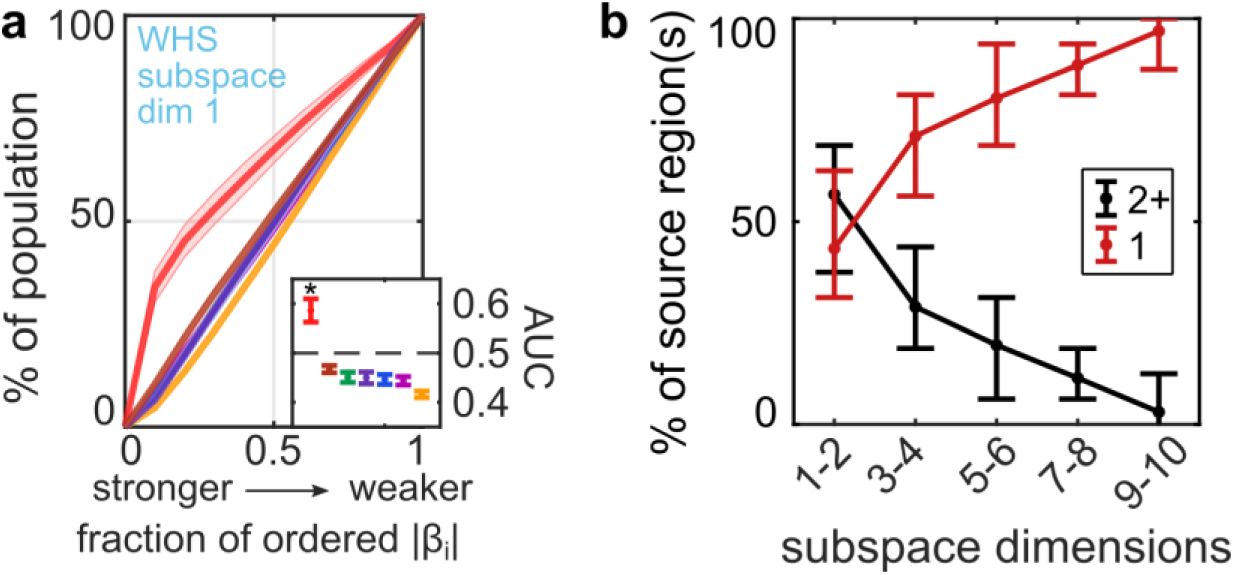
Subspace dimensions could integrate across multiple regions or reflect a ‘channel’ between two regions. Many subspace dimensions were functionally connected to a network of regions (Fig. 2). **(a)** However, as shown here, some subspace dimensions were dominated by a single source region. This can be thought of as creating a ‘channel’ between the source and target region. Panel shows the cumulative distribution of the percent of neurons within each source area contributing to the β weights along the first dimension of the whisker subspace, ordered from strongest-to-weakest. Presentation follows main Figure 2C; lines show mean, shaded region shows bootstrapped 95% confidence interval from n=70 datasets. Inset shows the mean and confidence intervals of the area under the curve (AUC) for the contributions from each region. Asterix denotes significance at p<0.05 versus an AUC of 0.5, bootstrap test. **(b)** Comparison of the percentage of subspace dimensions across regions that are functionally connected with two (or more) other regions (black) or a single region (red), split by subspace dimensions (x-axis). Line and error bars reflect mean and 95% confidence interval, respectively. While early subspace dimensions tended to be more integrative, later dimensions tended to be more channel-like.

**Supplementary Figure 7:**
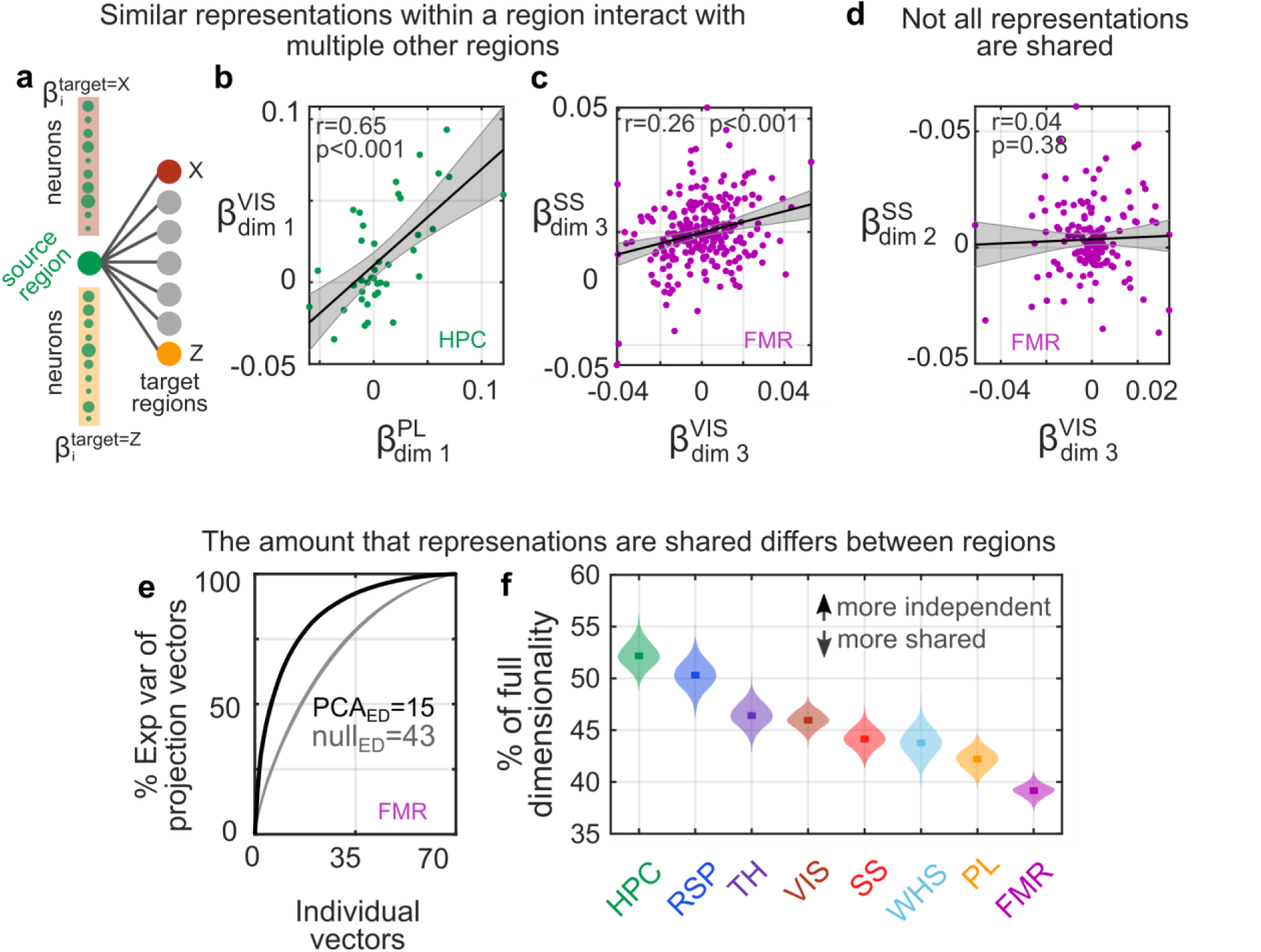
One representation within a region could project to multiple other regions. **(a)** Schematic showing how we estimated the vector of neural activity within a brain region that predicted activity in different target regions. **(b-c)** Example correlations of the projection vectors in **(b)** HPC and **(c)** FMR that project to other target regions. Superscript of X and Y axes show the target regions. Subscripts indicate the subspace dimensions of the target region. Dots show β weights of all neurons within HPC or FMR for an example dataset. p-values estimated with bootstrap (n=1000). **(d)** Example of independent (uncorrelated) projection vectors within FMR projecting to SS and VIS regions. Follows panels b and c. **(e)** Representative example dataset showing that projection vectors for FMR to other regions (black line) were more similar than expected by chance (gray lines; n=1000 area-label shuffles). X axis shows the total number of individual projection vectors (n=70; 7 target regions x 10 subspace dimensions per region). Y axis shows the cumulative percent explained variance of the projection vectors captured by their principal components. Text inset shows the effective dimensionality of the principal components for the original and shuffled data (see Methods). **(f)** Bootstrapped distribution showing the effective dimensionality of the projection vectors for each brain region relative to the full dimensionality (estimated from shuffled data, i.e., TrueED/NullED). Smaller percentages indicate regions with greater similarity in their projection vectors to other regions. In other words, the greater the percentage, the more independent (channel-like) each projection vector; the small the percentage, the more shared the projection vectors.

**Supplementary Figure 8.**
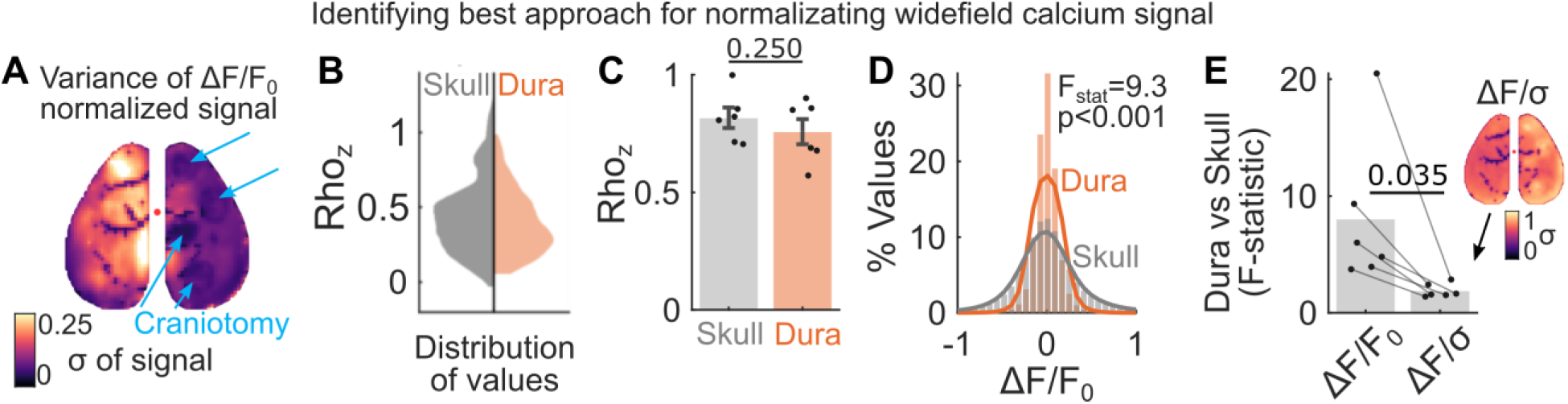
Optimizing the normalization of the widefield fluorescence signal. **(a)** Standard deviation of the fluorescence signal across cortical regions in an example recording. The activity of each pixel was normalized as the change in fluorescence over time relative to a rolling 120 second baseline activity (i.e., ΔF/F0). Left and right hemispheres show variance through intact skull and craniotomies, respectively. This reveals obvious differences in the variance of the fluorescence signal both across the cranium and between intact skull and craniotomies. Blue arrows indicate location of the craniotomies. **(b)** Differences in variance were not due to difference in patterns of activity, but rather signal magnitude (necessitating correcting for these differences). Violin plot shows the distribution of correlations in neural activity between corresponding pixels in each hemisphere in the example recording used in panel A. Left (gray) shows the correlation between the neural signal captured by pixels outside craniotomies (i.e., through the skull) on the left hemisphere with the same location (also through the skull) in the contralateral hemisphere. Right (orange) shows the correlation in neural activity between pixels within craniotomies (i.e., through dura and artificial Duragel) and their corresponding pixels on the contralateral hemisphere. Full distribution shown. **(c)** Patterns of activity within and outside craniotomies were similar in all recordings (n=6). Y-axis shows correlation between activity of through-skull pixels and dura-pixels with the contralateral hemisphere (e.g., as in B). p-value estimated by two-sided paired-permutation test. **(d)** While patterns of activity were similar, the magnitude of the signal differed between dura-pixels and skull-pixels. Histograms show distributions of pixel values from an example recording. The distribution of values of dura-pixels were significantly smaller than through-skull pixels. A potential source of this difference could be that removal of the skull leads to less scattering^54^, and in turn, fewer neurons contributing to the pixels within a craniotomy. p-value estimated with F-test of equal variances. **(e)** Normalization of fluorescence using the rolling 120 second standard deviation of each pixel (i.e., ΔF/σF) significantly reduced differences in signal magnitude between dura-pixels and skull-pixels. Points show the F-statistic comparing dura and skull-pixel distributions using the two normalization methods. p-value estimated with two-sided paired permutation test. Inset shows variance across imaged pixels (as in panel a).

**Supplementary Figure 9.**
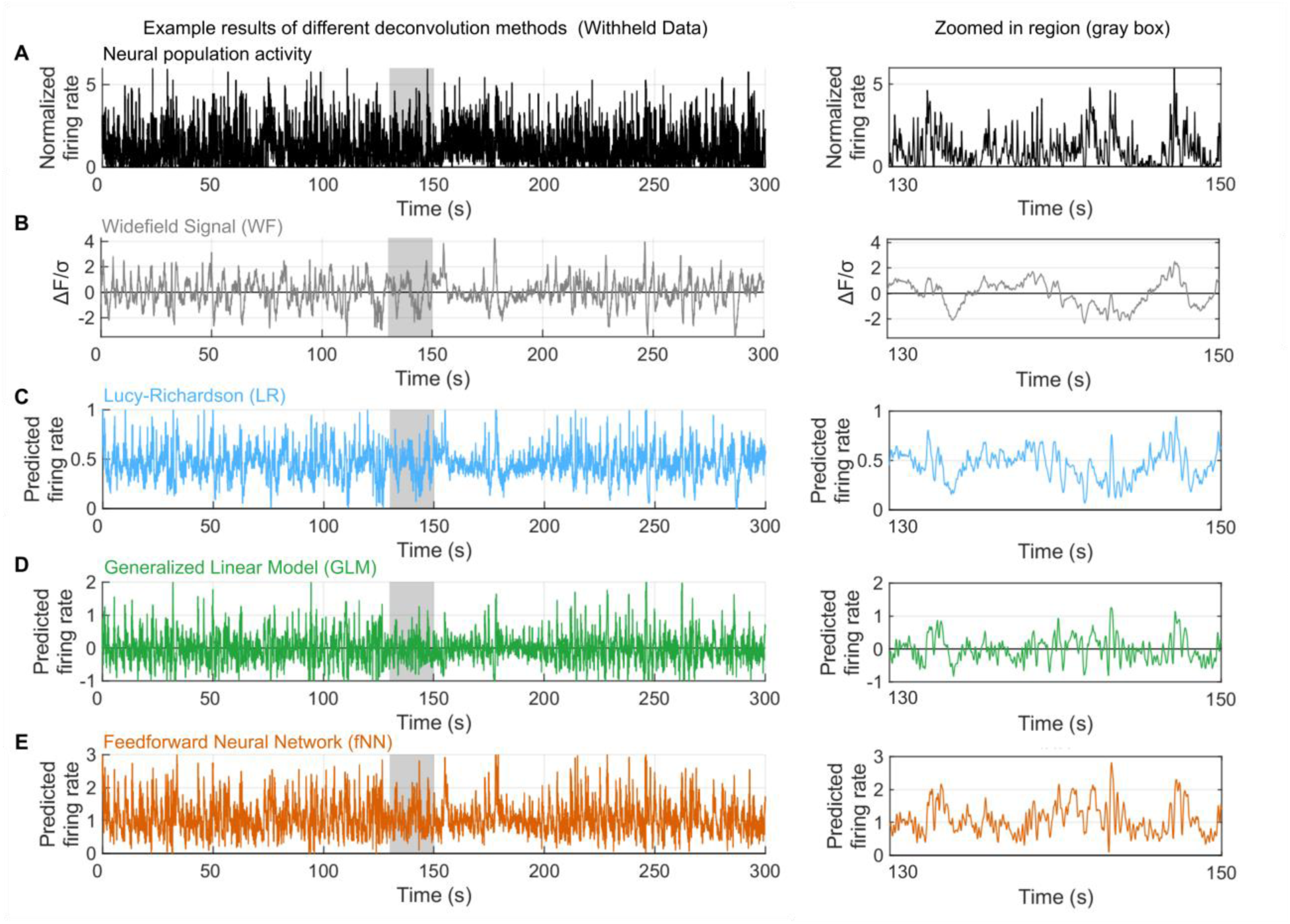
Comparison of methods for estimating neural population activity from widefield fluorescence signal. **(a)** Example ground truth spiking activity of neurons in the top 600uM of retrosplenial cortex. For each neuron, firing rate was normalized by the standard deviation across 90 minutes recording. Traces reflect mean firing rate of n=87 recorded neurons. Right inset shows zoomed in 20 second period (gray shaded region). **(b)** Normalized fluorescence signal from widefield imaging averaged across a 2-pixel radius surrounding the electrophysiologically recorded neurons in panel a. **(c-e)** Example results from three deconvolution approaches. **(c)** Lucy-Richardson (LR) deconvolution estimates spiking activity from fluorescence signal using a model of GCaMP6f kinetics. This approach has been used previously to estimate neural activity underlying the widefield imaging signal^5,38,39^. Methods for LR deconvolution followed previous work^38^ but with decay parameter γ = 0.89, which initial analyses found to be optimal for correlating to ground truth spiking activity. **(d)** Generalized linear modeling identifies a linear deconvolution kernel that best predicts firing rate from known spiking activity (methods followed previous work^37^). The deconvolution kernel was fit using the one-second of fluorescence signal on either side of the predicted timepoint. **(e)** Nonlinear deconvolution using a feedforward neural network. Just as with the GLM, the network was trained to predict spiking activity using the one-second of fluorescence on either side of the predicted timepoint. The network consisted of a single 20-neuron hidden layer with a tansig input function and positive linear output function. Model fitting for panels d and e were performed on a separate dataset and thus results show generalized performance to withheld data.

**Supplementary Figure 10.**
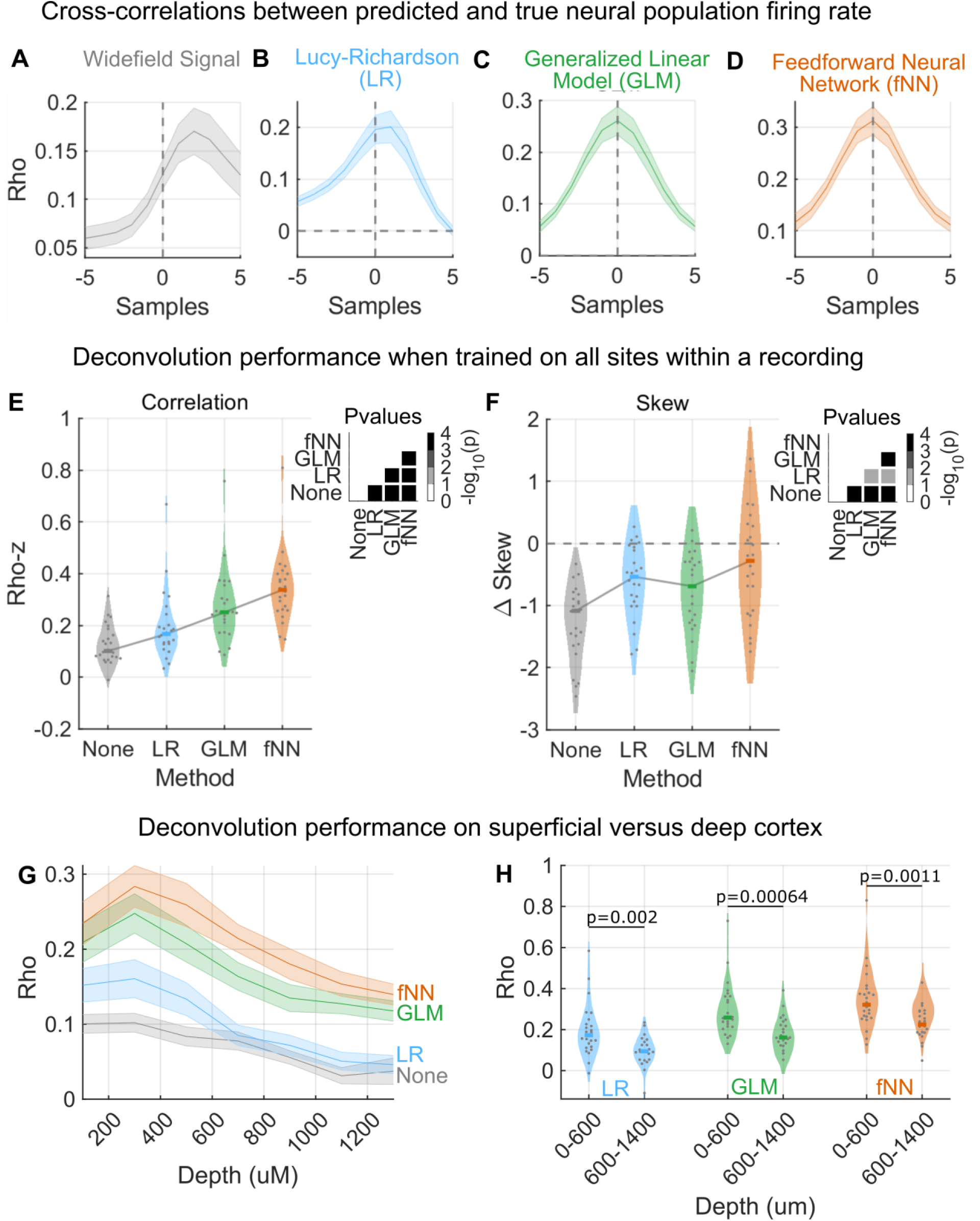
Identifying optimal deconvolution approach for estimating neural population spiking activity. **(a-d)** Cross correlation between the firing rate predicted from fluorescence signal and the true firing rate recorded by electrophysiology. **(a)** As expected, raw fluorescence was delayed relative to underlying spiking activity. **(b)** Lucy-Richardson deconvolution improved but did not fully resolve this delay. **(c)** Generalized linear models and **(d)** feedforward neural networks resolved the delay. Line and shaded region show mean and SE, respectively of n=24 comparisons on withheld data (from n=4 cortical recording sites, n=6 recordings). Deconvolution performed using activity from neurons in top 600uM of cortex. **(e)** Correlation between predicted spiking activity and ground truth spiking activity using different deconvolution methods. Full data distribution shown (n=24). Prediction was performed using fNN and GLM models fit to data from all four cortical sites for each recording. Y-axis shows R-to-Z transformed correlation coefficients. Inset shows pairwise statistical comparisons between methods (one-tailed, Signed-Rank Test; not corrected for multiple comparison). **(f)** Difference in skew in the distribution of predicted versus recorded firing rates over time within each recording. Neural spiking activity is typically log-normally distributed (i.e., highly skewed). Although this was not explicitly part of the fitting procedure, an optimal deconvolution method should estimate the skew observed in neural activity (i.e., Δ Skew should be close to zero). Negative values mean that the deconvolution method was less skewed than recorded spiking activity. fNN best recovered underlying spiking statistics compared to other methods. For main text experiments, fluorescence signal was deconvolved using networks trained on all areas (as shown here) from the entire recording. **(g)** Correlation between predicted spiking activity and ground truth spiking activity at different depths. Depths were sampled in 200uM windows, each overlapping by 100uM **(h)** Regardless of deconvolution method, fluorescence in the Thy1-GCaMP6f line used in this study best correlated with spiking activity in superficial (0-600uM) cortical layers. Full distribution (n=24) shown, p-values estimated with one-tailed signed-rank test.

**Supplementary Figure 11.**
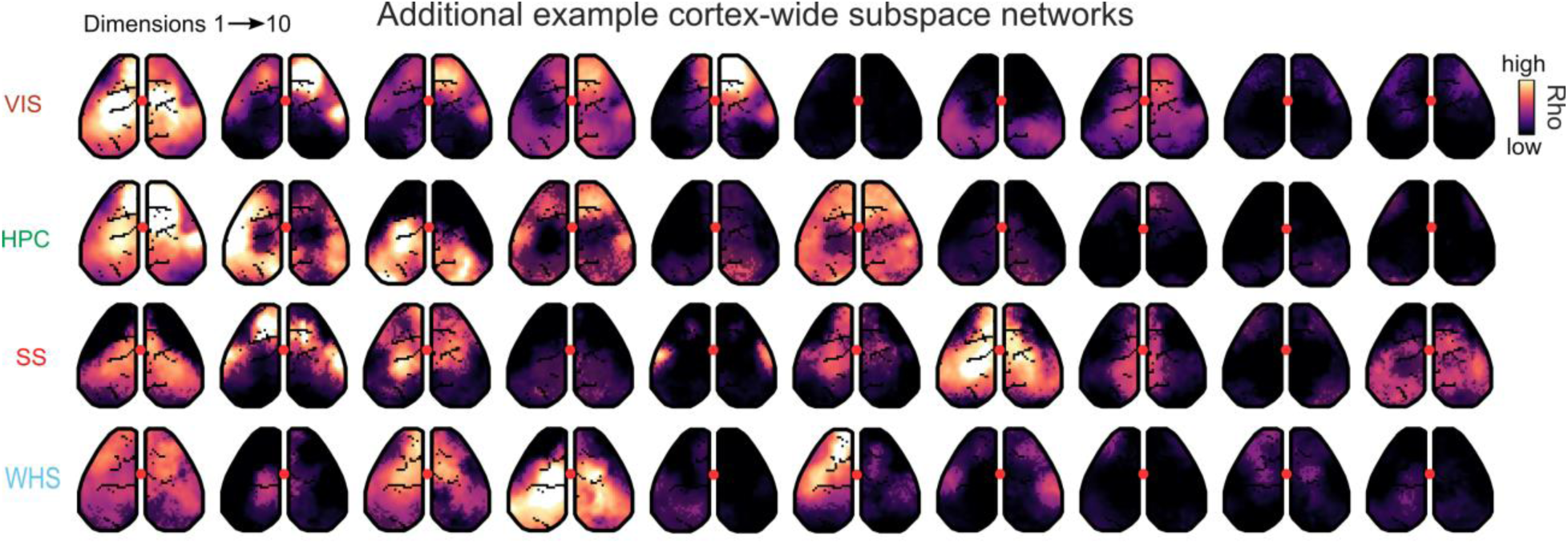
Additional examples of cortex-wide subspace networks. Presentation follows Figure 3b. Maps in each row are ordered by their subspace dimension of subspaces for a target region.

**Supplementary Figure 12.**
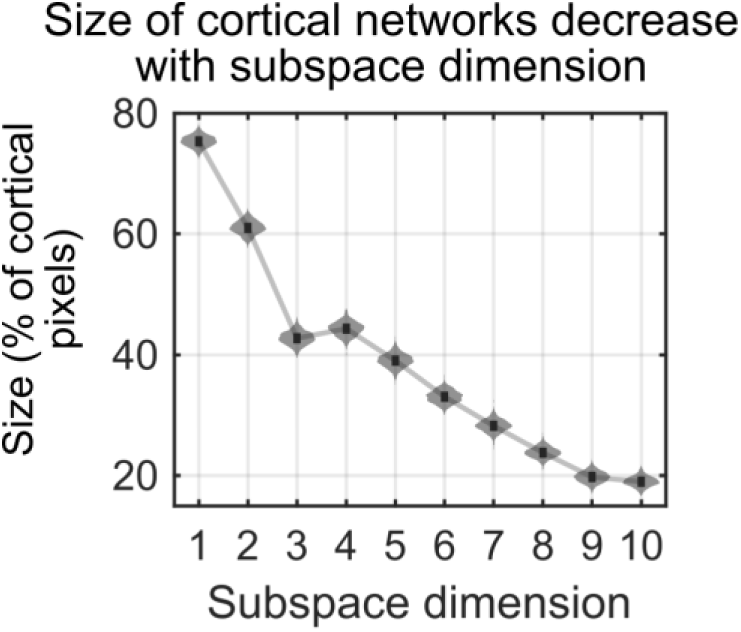
Early subspace dimensions reflect a broader cortex-wide subspace networks. Bootstrapped distribution showing the percentage of the cortex (y-axis) engaged as part of the cortical subspace network of all subspace dimensions (x-axis). For example, the cortical network associated with subspace dimension 1 was significantly larger than subsequent dimensions. Subspace dimension 2 was also significantly larger than subsequent dimensions (p<0.001, n=1000 bootstraps from n=630 datasets).

**Supplementary Figure 13.**
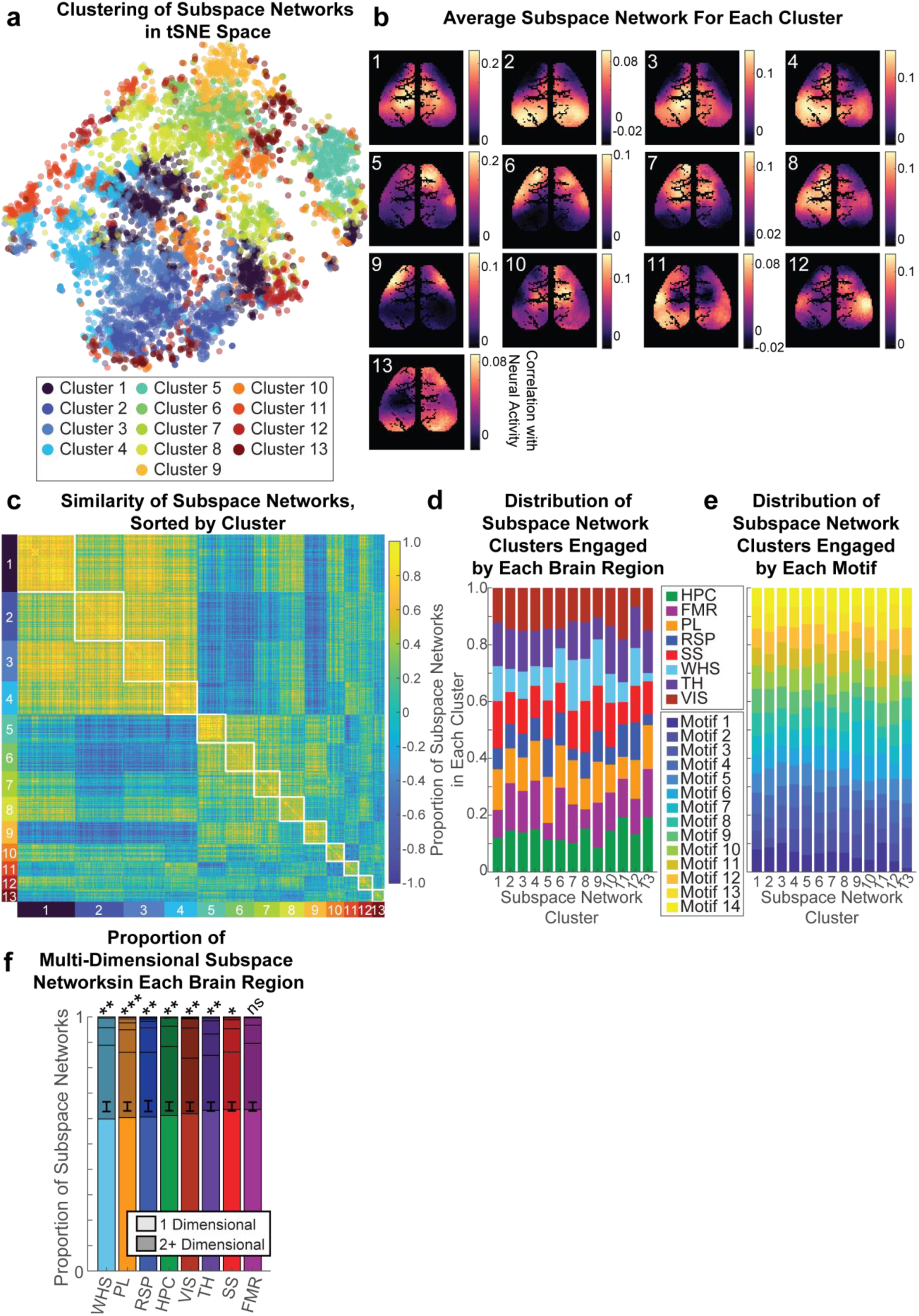
Clustering of subspace dimensions. **(a)** Distribution of subspace networks projected into tSNE space. Distance is the inverse correlation. Each point reflects a subspace network associated with a different dimension, across all areas, dimensions, and datasets. Points are colored by their assigned cluster. Points are partially transparent to facilitate seeing all data points. **(b)** Median subspace network for each cluster. Color axis shows strength of correlation. Only pixels that were valid across all recordings were used (i.e., the map of vasculature is combined across all recordings). Clusters were labeled according to the number of subspace networks in each cluster (descending). **(c)** Similarity matrix for all subspace networks, sorted by cluster identity. Cluster groupings are shown along the x and y axes. Colorbar shows correlation of subspace network across the entire spatial map. White boxes indicate the within-cluster pairs of subspace networks. **(d)** Proportion of subspace networks (y-axis) in each cluster (x-axis) that was associated with each brain region (color label). All subspace networks were seen in all brain regions. **(e)** Similar to panel d, but now showing the proportion of subspace networks observed in each motif. **(f)** Proportion of subspace networks (y-axis) that engage one (light colors) or more (dark colors) dimensions within a brain region. Subspace networks were considered to be the same if they fell into the same cluster. Vertically stacked sub-divisions of bar indicate the proportion of subspace networks associated with one (light colors), two, three, etc. dimensions (dark colors). Proportions are averaged across all datasets within the same region. Black errorbars show mean and standard error of the expected proportion, given the overall distribution of clusters. Significance was measured with a permutation test. *** p<0.001, ** p<0.01, * p<0.05. Seven of the eight regions showed a significantly greater number of multi-dimensional subspace networks than expected by chance. Across all regions, the proportion of multi-dimensional subspaces was significant (p=2e-5).

**Supplementary Figure 14.**
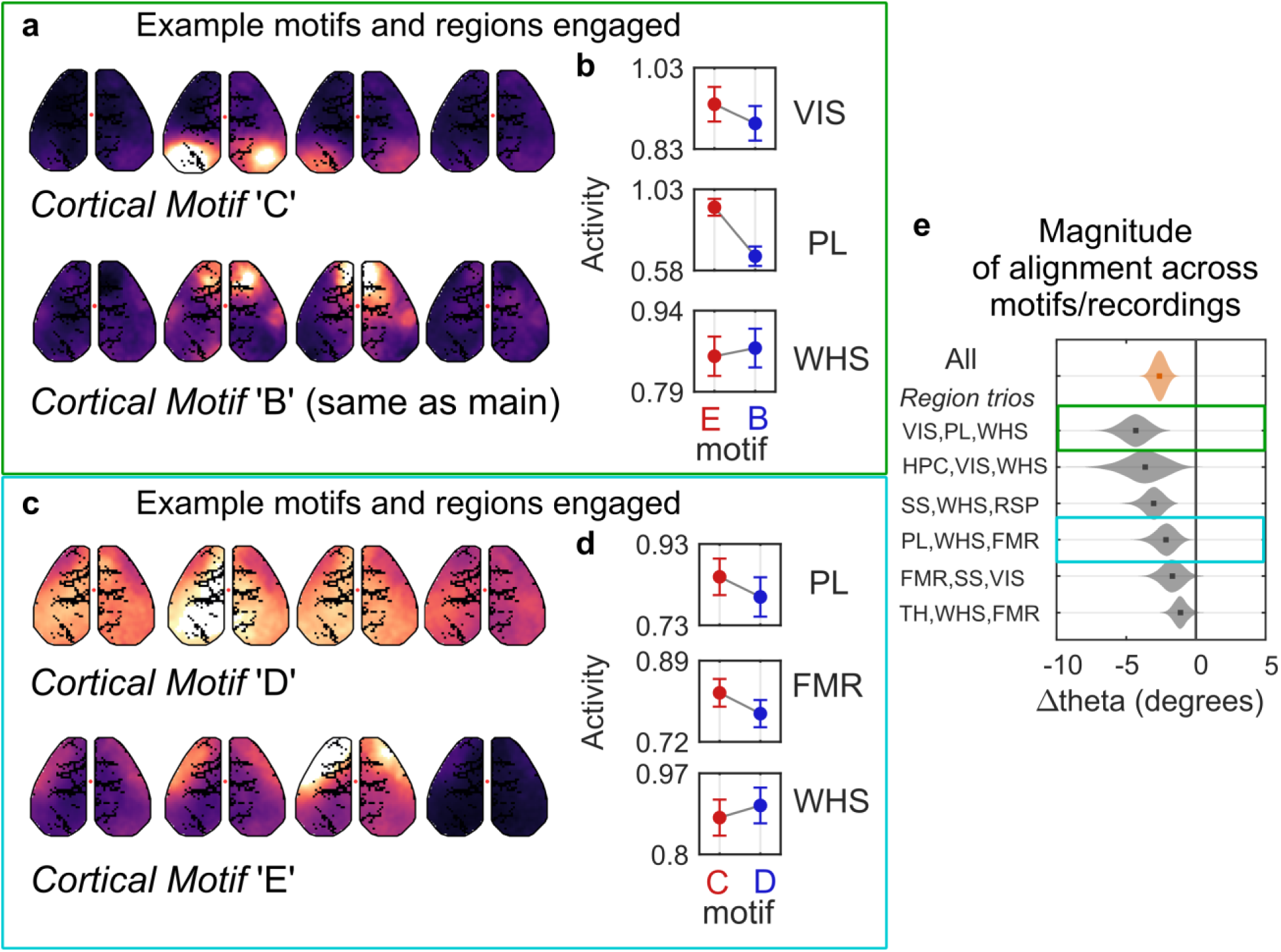
Additional examples of compared pairs of motifs and trios of brain regions for analysis of alignment between population representations and subspace activity. Pairs of motifs and trios of brain regions were selected to test what may underlie differences in the flow of neural activity between areas (see Methods for selection criteria and blinding). **(a-b)** Comparison of activity between motifs C and B showing that they engage Visual, Prelimbic, and Whisker regions but in unique ways. **(c-d)** Comparison of activity between motifs D and E showing that they engage Prelimbic, Frontal Motor, and Whisker regions but in unique ways and differ in their alignment angles. **(e)** Bootstrapped distribution of difference in alignment from all pairs of motifs and trios of brain regions across recordings (replicated from main Fig. 6c). Boxes outline the distributions that correspond to the examples in panels a-d).

**Supplementary Figure 15.**
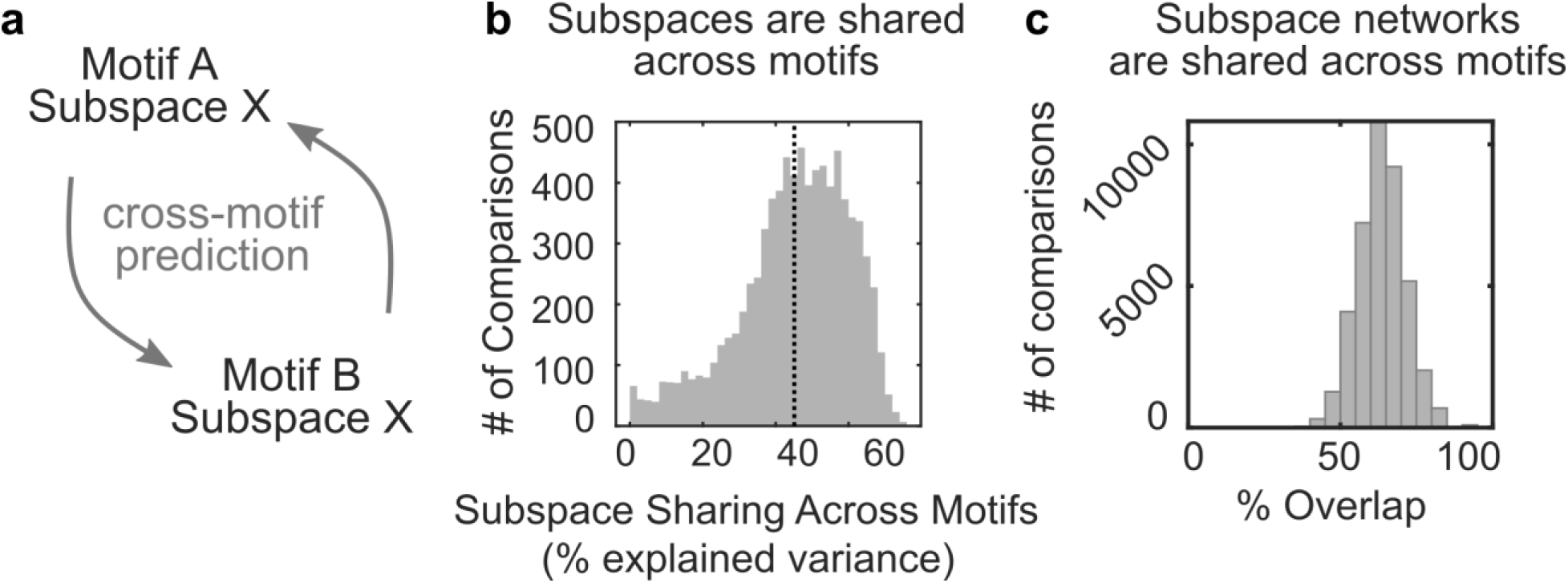
Subspaces networks were stable across motifs. **(a)** Independent regression models fit during one motif were used to predict the interregional subspace activity during another motif. **(b)** Histogram showing the distribution in explained variance captured by cross-motif predictions, across all pairs of motifs and all regions. Dotted line shows mean. **(c)** Histogram of similarity in cortical networks maps across different motifs, taken as the percent of overlapping significant pixels. Data reflect the best match between a map for one dimension/source region for a motif and any of the first ten dimensions of that same source region in a different motif (to adjust for any reordering of dimensions across motifs).

